# Host MOSPD2 enrichment at the parasitophorous vacuole membrane varies between *Toxoplasma* strains and involves complex interactions

**DOI:** 10.1101/2022.12.23.521848

**Authors:** Abel Ferrel, Julia Romano, Michael W. Panas, Isabelle Coppens, John C. Boothroyd

## Abstract

*Toxoplasma gondii* is an obligate, intracellular parasite capable of causing severe disease in warm-blooded animals. Infection of a cell produces a unique niche for the parasite named the parasitophorous vacuole (PV) initially composed of host plasma membrane invaginated during invasion. The PV and its membrane (PVM) are subsequently decorated with a variety of parasite proteins allowing the parasite to optimally grow in addition to manipulate host processes. Recently, we reported a proximity-labeling screen at the PVM-host interface and identified host ER-resident MOSPD2 as being enriched at this location. Here we extend these findings in several important respects. First, we show that the extent and pattern of host MOSPD2 association with the PVM differs dramatically in cells infected with different strains of *Toxoplasma*. Second, in cells infected with Type I RH strain, the MOSPD2 staining is mutually exclusive with regions of the PVM that associate with mitochondria. Third, immunoprecipitation and LC-MS/MS with epitope-tagged MOSPD2-expressing host cells reveals strong enrichment of several PVM-localized parasite proteins, although none appear to play an essential role in MOSPD2 association. Lastly, most MOSPD2 associating with the PVM is newly translated after infection of the cell and requires the major functional domains of MOSPD2, identified as the CRAL/TRIO domain and tail anchor, although these domains were not sufficient for PVM association. Collectively, these studies provide new insight into the molecular interactions involving MOSPD2 at the dynamic interface between the PVM and the host cytosol.

**Importance:** *Toxoplasma gondii* is an intracellular pathogen that lives within a membranous vacuole inside of its host cell. This vacuole is decorated by a variety of parasite proteins that allow it to defend against host attack, acquire nutrients, and interact with the host cell. Recent work identified and validated host proteins enriched at this host-pathogen interface. Here, we follow up on one candidate named MOSPD2 shown to be enriched at the vacuolar membrane and describe it as having a dynamic interaction at this location depending on a variety of factors. Some of these include the presence of host mitochondria, intrinsic domains of the host protein, and whether translation is active. Importantly, we show that MOSPD2 enrichment at the vacuole membrane differs between strains indicating active involvement of the parasite with this phenotype. Altogether, these results shed light on the mechanism and role of protein associations in the host-pathogen interaction.

## Introduction

*Toxoplasma gondii* is an obligate, intracellular parasite in the Apicomplexa phylum. It is estimated that one in three people globally is infected. Although most cases are mild or asymptomatic, severe disease can occur in individuals that are immunocompromised or in congenitally infected fetuses [1]. The symptomatic stage of infection is caused by a fast-replicating form called tachyzoite.

Tachyzoites, like the other stages of *Toxoplasma’s* life cycle, have three unique sets of organelles that allow them to invade a host cell and set up a niche for intracellular growth. Preceding invasion, the first set of these organelles called micronemes are deployed for the main role of attachment [2], [3]. Rhoptries are the second set of invasion organelles to be deployed and their protein contents can be subdivided into two main classes, the rhoptry neck (RON) proteins that aid in invasion and the rhoptry bulb (ROP) proteins that can co-opt host functions [4]. After the deployment of ROPs, a tachyzoite invades the host cell by mechanically pulling itself in and invaginating the host plasma membrane to form the PV with its delimiting membrane, the PVM [5]–[10]. Remarkably, this process sieves many host proteins from the PVM which grants it the privilege of being non-fusogenic with harmful host lysosomes or endosomes [11]. During and after the formation of the PV, the parasite secretes proteins from its third set of unique organelles termed dense granules with their protein contents termed GRAs [12]. Characterized GRAs have myriad functions including nutrient acquisition [13], interactions with mitochondria [14], and manipulation of host signaling pathways [15], [16].

The PVM is a dynamic interface that separates the host cell and parasite. Association of host organelles derived from the exocytic pathway (e.g., endoplasmic reticulum and Golgi apparatus), endocytic pathway (endolysosomes and lysosomes), and mitochondria has been described to occur at the PVM [17]–[19]. Host mitochondrial association (HMA) is a strain-dependent phenotype mediated by the parasite’s mitochondrial association factor 1b (MAF1b) found at the PVM [14], [20]; Type I and III parasites express MAF1b and recruit mitochondria, while Type II parasites do not express the protein and therefore do not display HMA [20]. Interestingly, engineering expression of MAF1b in Type II parasites is sufficient for HMA to occur [14]. Although the parasite effector mediating HMA has been described, the parasite proteins that allow association of other organelles have remained elusive.

In contrast to proteins derived from the parasite, only a few host proteins have been described to localize to the PVM post-invasion. Of the few, host Immunity Related Guanosine Triphosphatases (IRGs), along with p65 guanylate binding proteins (GBPs), are encoded by IFN-γ-stimulated genes involved in cell-autonomous immune defense against *Toxoplasma* in rodents [21], [22]. They localize to the PVM and lead to permeabilization of the vacuolar membrane, allowing clearance of the parasite. To combat this, *Toxoplasma* injects ROP5/17/18 during rhoptry discharge and these eventually associate with the host-cytosolic face of the PVM [23]– [25]. These are all ROP2-family members with arginine-rich amphipathic helix (RAH) domains that mediate PVM association [26]. Functionally, ROP17 and ROP18 are serine/threonine kinases that inactivate host IRGs by phosphorylating them [23], [25]. ROP5 is a catalytically inactive pseudokinase that binds IRGs, altering their shape and making them available for ROP18 inactivation [24]. Because of this crucial role, polymorphisms in ROP5/17/18 sequences and/or expression levels are associated with dramatic differences in virulence in mice, at least [27]–[31]. More recently, two PVM-localized GRAs, GRA14 and GRA64, have been shown to interact with host Endosomal Sorting Complex Required for Transport (ESCRT) subsequently leading to internalization of proteins from the host cell [32], [33].

Recently, Cygan et al. performed an unbiased proteomic screen to identify additional host and parasite PVM proteins by localizing the promiscuous biotin ligase, miniTurbo, to the host-cytosolic face of the PVM [34]. Among the host proteins detected as enriched at the PVM were accessory proteins to the ESCRT complex. Additionally, and unexpectedly, the ER-anchored membrane protein motile sperm domain-containing protein 2 (MOSPD2) was also identified and subsequently validated to be at the PVM using a transiently expressed, epitope-tagged construct [34]. MOSPD2 has been described in prior studies of uninfected cells as mediating inter-organellar associations through protein-protein interactions via its Major Sperm Protein (MSP) and cellular retinaldehyde-binding protein (CRAL)/triple functional domain protein (TRIO) domain [35], [36]. It was tantalizing to hypothesize, therefore, that MOSPD2 might be mediating host ER-PVM interactions in infected cells, and in this study, we utilize microscopy, genetic, and biochemical techniques to investigate the mechanism and function of MOSPD2 localization at the PVM. We take advantage of three common strains of *Toxoplasma* to narrow down possible functions. In addition, biochemical methods allowed us to immunoprecipitate MOSPD2 and identify interacting *Toxoplasma* proteins at the PVM which were then tested for a role in and/or dependence on MOSPD2 association. We also assessed MOSPD2 association in the context of other parasite/host interactions at the PVM. The results reveal a unique interaction between *Toxoplasma* tachyzoites and the host cells they infect.

## Results

Association of MOSPD2 with the PVM was previously described by Cygan et al. using Type I RH. To determine whether MOSPD2 association differs between strains of *Toxoplasma*, human foreskin fibroblasts (HFFs) were infected with either Type I RH, Type II ME49, Type III CTG, or the close relative apicomplexan parasite, *Neospora caninum* (Nc) for 21 hours and then stained using anti-MOSPD2 antibodies. The results showed association of MOSPD2 with the PVMs of cells infected with RH and ME49 at substantially higher levels than those infected with CTG and Nc (Fig. 1A). To quantify these apparent differences, we used Fiji to measure the florescence intensity at the PVM 21 hours post infection (hpi) while excluding the host cytoplasm and lumen of the PV. The results showed that while there was considerable variability between the mean fluorescence intensity of each PVM in a given monolayer, overall, the PVMs in cells infected with RH and ME49 averaged almost twice the fluorescence intensity of cells infected with CTG or *Nc* (Fig. 1B). To more precisely characterize this association, we performed immune-electron microscopy (IEM) with MOSPD2 antibodies on RH-infected HFFs. IEM images showed MOSPD2 located at the ER-PVM interface, although the extremely close apposition of these two membranes does not allow determination of precisely which membrane MOSPD2 is anchored within (Fig. 1C).

**Fig 1:**
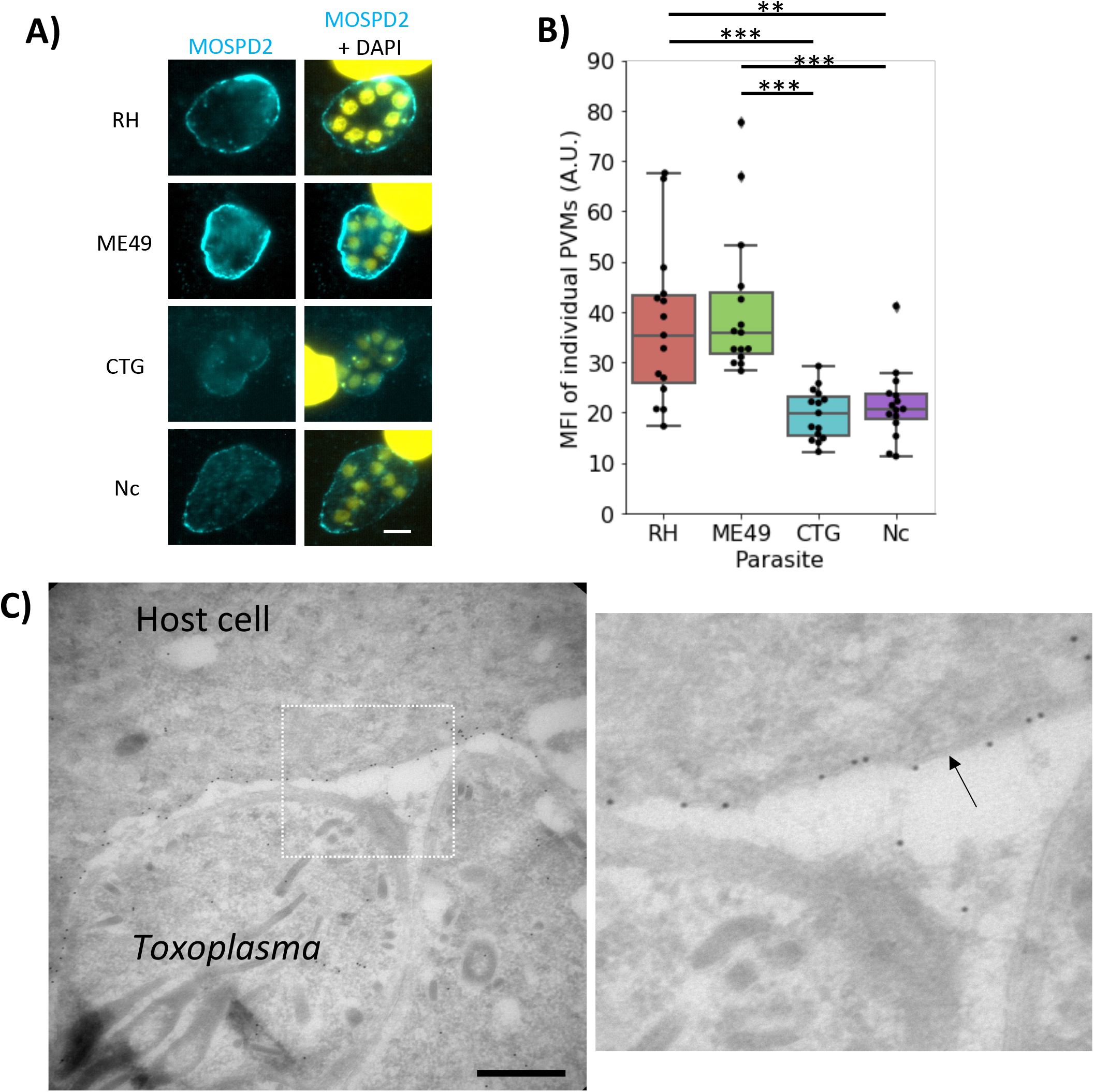
MOSPD2 localizes to the PVM and differs between strains. A. Representative images of HFFs infected with either RH, ME49, CTG, or Neospora caninum (Nc) for 21 h prior to fixing with methanol. Monolayers were then stained for endogenous MOSPD2 (cyan) and DAPI (yellow). Scale bar = 5 μm. B. Bar plot showing mean fluorescence intensity (MFI) of MOSPD2 signal on the PVM of individual vacuoles 21 hpi. Whiskers show the maximum value within 1.5 of the interquartile range. Data are from one experiment that pooled three biological replicates. One of three fully independent experiments is shown. Significance was tested using a One-way ANOVA and Tukey Post-HOC test thereafter. (** indicates P<0.01, *** indicates P<0.001). C. Left panel, electron micrograph of 24 h RH-infected HFFs stained with MOSPD2 antibody (1/50) and visualized using Protein A-gold particles. Scale bar = 500 nm. Right panel, zoomed image of boxed region in left image with an arrow pointing at the PVM.

Although the major difference in MOSPD2 association seen during infection is between RH/ME49 and CTG/*Nc*, RH and ME49 also appear to differ in the pattern of MOSPD2 association at the PVM (Fig. 1A). To quantify the fluctuation in MOSPD2 signal in infected cells, the PVM-localized fluorescent signal was measured using Fiji (see methods). MOSPD2 fluorescence at the PVM appeared considerably more patchy (i.e. had more variation in fluorescence intensity around any given PVM) in cells infected with RH than ME49 (Fig. 2A and B). A major difference known for the PVM of cells infected with RH vs. ME49 is the phenomenon of host mitochondrial association or HMA [14]. MOSPD2 is known to have a role in membrane contact sites between organelles and is present at ER-mitochondria interface in HeLa cells [35]. Therefore, it was possible that MOSPD2 might be playing a role in HMA at the PVM. To first determine whether MOSPD2 co-localizes with mitochondria at the PVM, RH-infected HFFs were stained with MitoTracker and antibodies to endogenous MOSPD2. The results showed that localization of host mitochondria and MOSPD2 at the PVM were, in fact, anti-correlated (Fig. 2C). This was confirmed by immuno-electron microscopy with the results showing MOSPD2 at the PVM only in regions that did not have HMA (Fig. 2D). These results suggest that MOSPD2 can only associate with regions of the PVM that are not occupied by host mitochondria.

**Fig 2:**
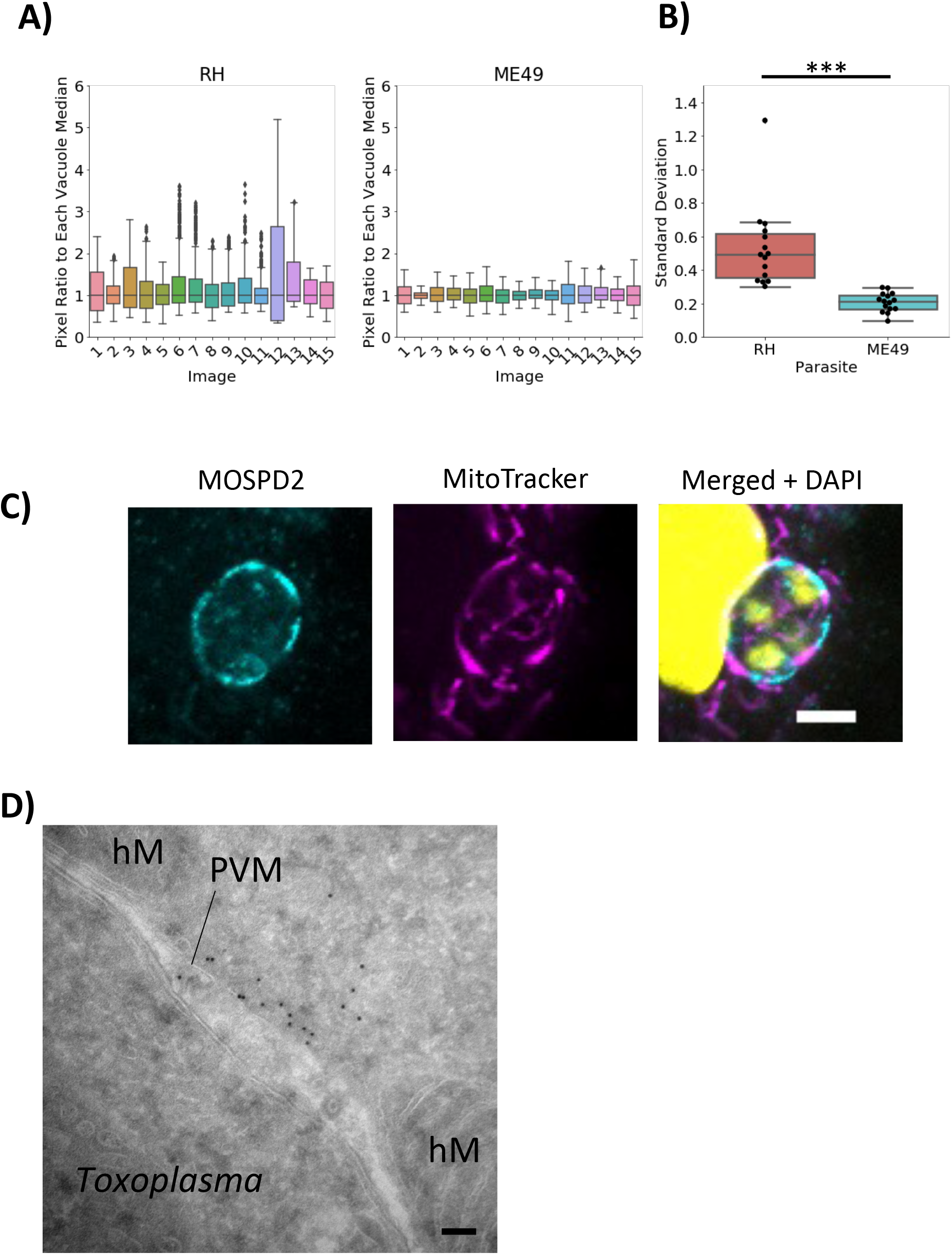
MOSPD2 associates at the PVM where host mitochondria are not recruited. A. HFFs were infected with RH or ME49 for 21-24 h, then fixed with methanol and stained for MOSPD2 and calreticulin. MOSPD2 fluorescence intensity at the PVM from 15 vacuoles in each strain were quantified (see methods). The fluorescence intensity at each pixel was divided by the median of that respective vacuole to normalize all vacuoles to the median (y-axis). Data are from one experiment that pooled three biological replicates. Data are representative of two independent experiments. B. The standard deviation (y-axis) of the ratios from (A) for each vacuole were plotted. Significance was tested using Student’s t test (*** indicates P<0.001). C. HFFs were infected with RH parasites for 21 h, stained with MitoTracker (see Methods), fixed with paraformaldehyde, and permeabilized with 0.1% Triton-X100. Stains include mitochondria (magenta), MOSPD2 (cyan), and DAPI (yellow). Data are representative of two independent experiments. Scale bar = 5 μm. D. Electron image of the PVM of RH-infected HFFs stained with MOSPD2 antibody and visualized with Protein A-gold particles thereafter. hM, host mitochondria. Scale bar = 100 nm.

The parasite protein responsible for HMA is known to be MAF1b [14]. It colocalizes with host mitochondria at the PVM and is not expressed in ME49. To determine if MAF1b influences the loci where MOSPD2 localizes, whether directly or indirectly, HFFs were infected with ME49 parasites that ectopically express MAF1b. Immunostaining showed wild type ME49 had low signal fluctuations around the PVM while ME49 that express MAF1b had MOSPD2 signal mirroring the pattern seen in RH; i.e., MOSPD2 appeared absent wherever MAF1b was present (Fig. 3 A-C). While we cannot exclude the possibility that MAF1b itself directly prevents MOSPD2 association, these results seem most likely to be due to active exclusion by host mitochondria recruited to the PVM.

**Fig 3:**
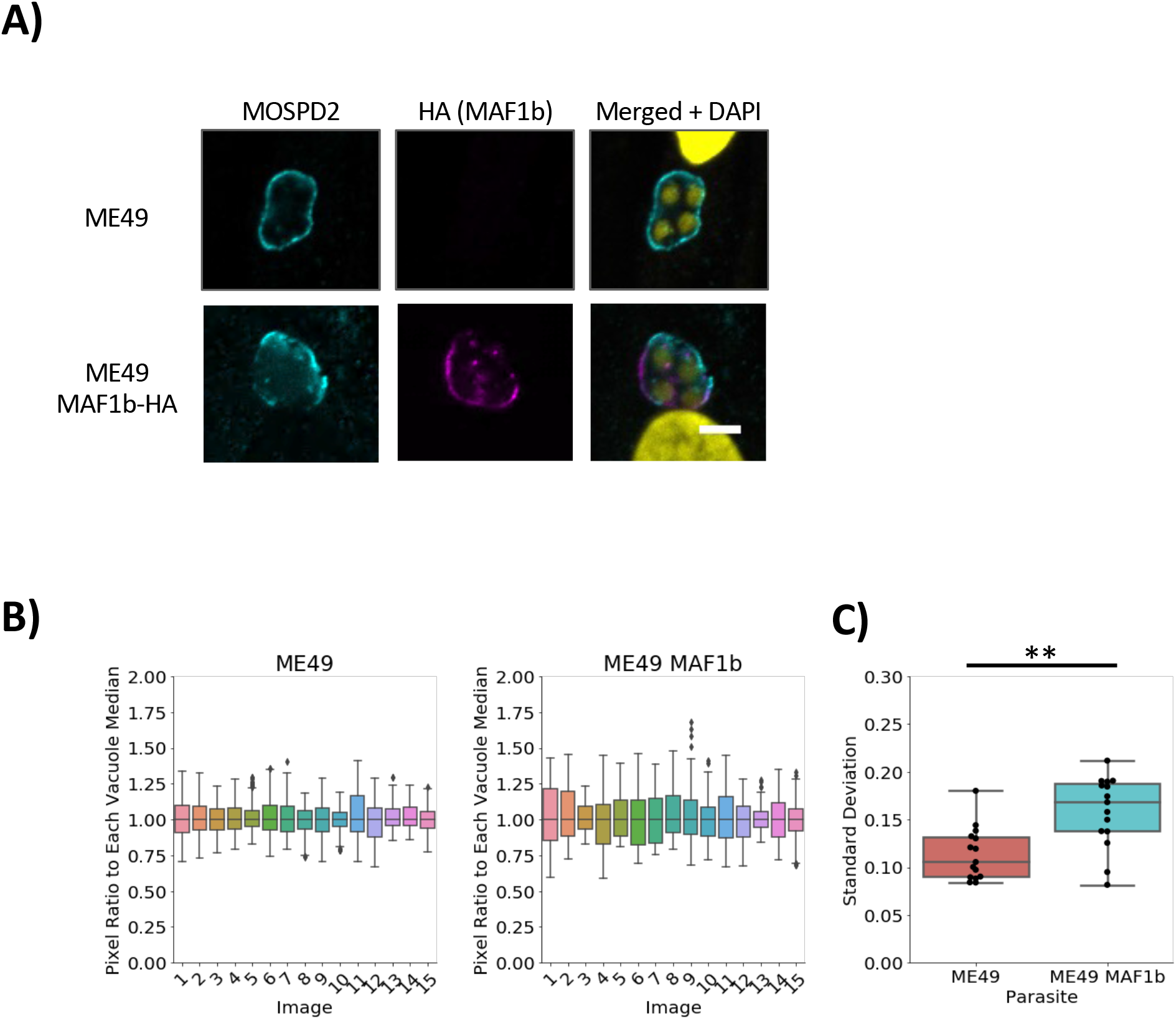
MOSPD2 association at the PVM is altered in ME49 by ectopically expressed MAF1b. HFFs were infected with either ME49 or ME49 MAF1b-HA for 21 h then fixed with methanol. A. Representative images from two independent experiments show stains for MOSPD2 (cyan), HA (MAF1b, magenta), DAPI (yellow). Scale bar = 5 μm. B. HFFs were infected with ME49 or ME49 MAF1b-HA as described. MOSPD2 normalization measurements were done as described in Fig. 2 and methods. Representative data are from one experiment that pooled three biological replicates. Data are representative of two independent experiments C. Standard deviations (y-axis) plotted for each strain from (B). Significance was tested using Student’s t test (** indicates P<0.01).

The so-called MYR complex is known to translocate soluble GRA effector proteins across the PVM and into the host cell [37]–[39]. The first component characterized of the MYR complex was MYR1 and knocking out this gene leads to the inability of GRA effectors (e. g., GRA16, GRA24, etc.) to translocate across the PVM and into the host cell. *Toxoplasma* effectors have a variety of functions and to explore whether one or more of the MYR1-mediated set might be responsible for the MOSPD2-association phenotype, HFFs infected with MYR1-knockout strains were assessed by immunofluorescence [40]. The results showed that the absence of *MYR1* in ME49 did not affect recruitment of MOSPD2 to the PVM (Fig. 4A), with fluorescence quantifications on ME49 and ME49Δ*myr1* PVMs showing no statistical difference between the two (Fig. 4B). These data indicate that MYR1 and the MYR1-dependent GRA effectors are not needed for the association of host MOSPD2 at the PVM. GRA45 is a protein known to be important for proper insertion of a different class of GRA proteins, ones that are integral to (i.e., spanning) the PVM [41]. To test if any such GRAs were necessary for MOSPD2 association with the PVM, HFFs were infected with parasites having a disrupted *GRA45* gene. Results showed no difference between control and Δ*gra45* parasites (Fig. 4C and D) indicating that PVM-integral GRAs are not required for the presence of MOSPD2 at the PVM.

**Fig 4:**
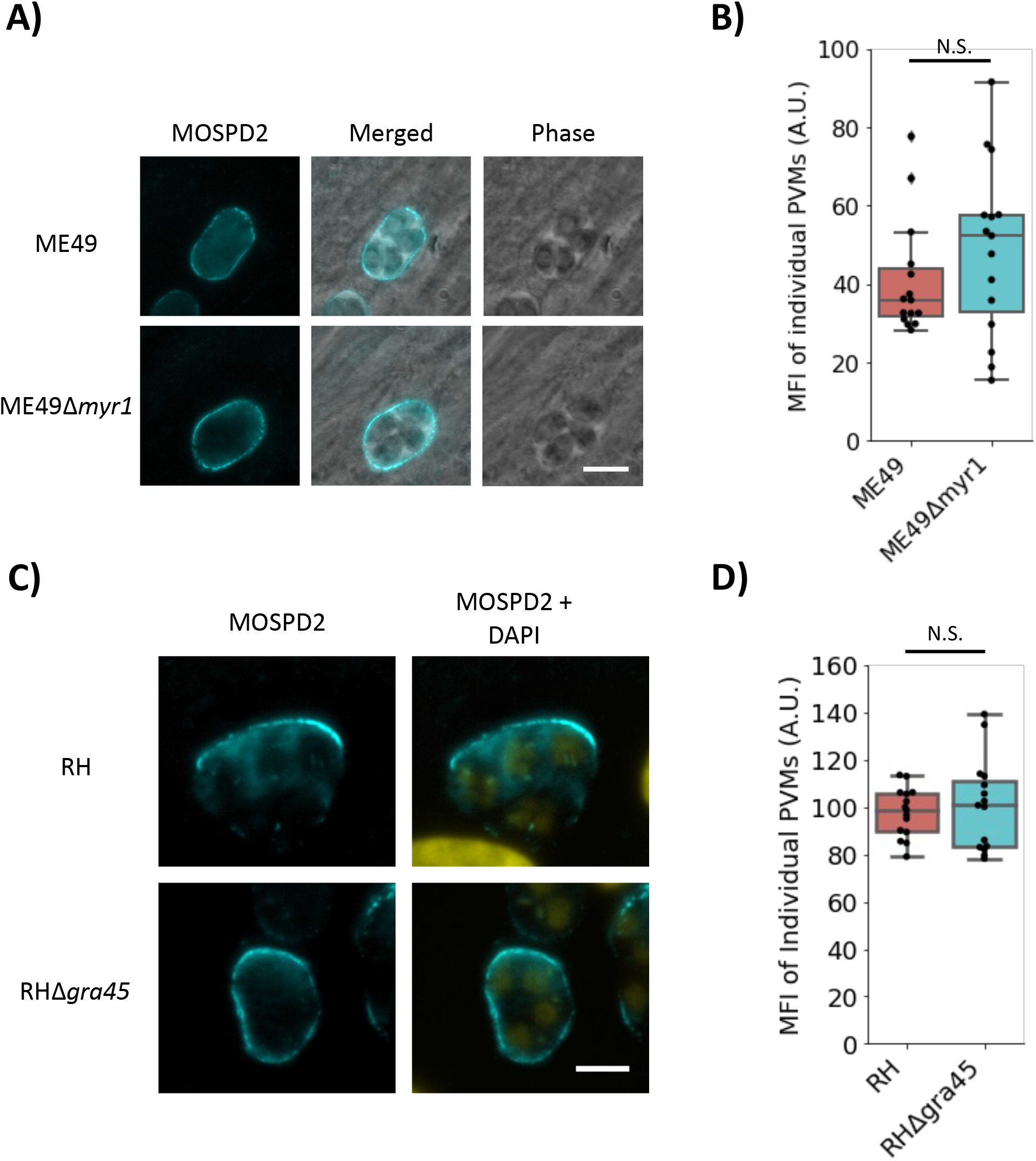
MYR1 and GRA45 are not necessary for host MOSPD2 association at the PVM. A. Representative images of ME49- or ME49Δmyr1-infected HFFs 21 hpi. Cells were then fixed with methanol and stained with anti-MOSPD2 antibodies. Scale bar = 10 μm. B. Boxplot of MOSPD2 fluorescence intensities at the PVM from vacuoles in (A). N.S. indicates not statistically significant (P≥0.05). Representative images (C) and quantitation (D) 21 hpi of HFFs infected with RHΔgra16::GRA16-HA parental (RH) and Δgra45 made in the same background (RHΔgra45). Samples were processed as described above. Scale bar = 5 μm. Quantitation was performed in Fiji as described in the methods. Data are from one representative of three independent experiments, each consisting of three pooled biological replicates. Significance was tested using Student’s t test for significance.

To further explore which *Toxoplasma* proteins at the PVM might interact with MOSPD2, HFFs overexpressing V5-tagged MOSPD2 were infected with ME49 and immunoprecipitated using anti-V5 nanobodies for LC-MS/MS identification. Western blot stains for V5 showed strong enrichment of the epitope-tagged protein in the eluted material (Fig. 5A). VAP-A, a host protein known to interact with MOSPD2 at the ER [42], was also enriched in the elution while calreticulin which is also an ER protein but is known to not associate with MOSPD2 was not co-precipitated and acted as a negative control (Fig. 5A). To determine the proteins most enriched in our immunoprecipitation, ratios between our V5-MOSPD2 and untagged control were generated (Fig 5B). The most enriched *Toxoplasma* protein was ROP17, a serine/threonine kinase known to be at the PVM and having at least two functions [25], [43]; and among the top 15 most enriched proteins, 7 (ROP1/5/17/18, GRA12/14 and MAF1a) have previously been reported to localize to the PVM [20], [25], [30], [44]–[48] indicating that our protocol was indeed enriching for proteins associating, directly or indirectly, with MOSPD2 at the PVM.

**Fig 5:**
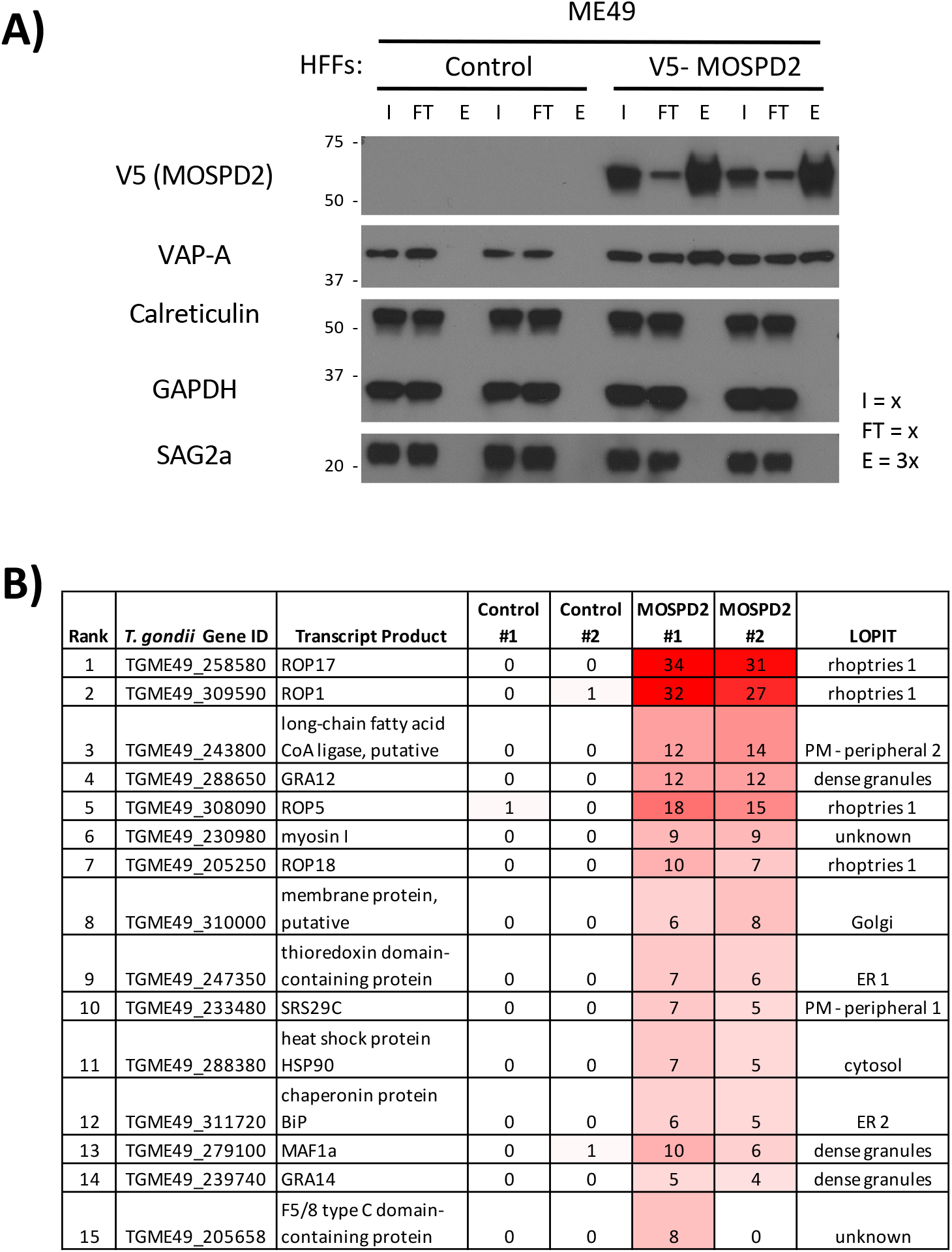
V5-MOSPD2 immunoprecipitation and LC-MS/MS identifies known T. gondii proteins at the PVM. A. ME49-infected control or V5-MOSPD2 expressing HFFs were infected for 19 h then lysed with RIPA. Whole cell lysates were incubated with anti-V5 nanobodies, spinning for 1 h at 4°C. Nonbinding lysates were collected, and beads were washed. Bound proteins were eluted in Laemmli buffer at 90°C for 5 minutes. Western blot was probed for V5 (MOSPD2), VAP-A, Calreticulin, GAPDH, and SAG2a. “x” values denote relative amount of the sample loaded into each lane (i.e., elution had three times as much of the total sample loaded relative to the input and flowthrough lanes) Data are representative of three independent experiments. B. Samples described in (A) were submitted for LC-MS/MS protein identification. A value of 1 was added to all spectral counts for control and V5-MOSPD2 conditions then averaged and the results for each parasite protein were listed in descending rank for enrichment in the MOSPD2 samples relative to controls [Supp. Data Set 1]. Intensity of red shading reflects relative number of spectral counts detected in each sample. LOPIT data describes subcellular localization within intact parasites for identified proteins [84]. Such data do not relate to a given protein’s final location within the infected host cell.

To determine whether the top *Toxoplasma* candidate proteins from the V5-MOSPD2 immunoprecipitation play a direct role in MOSPD2 association at the PVM, HFFs were infected with wild type parasites or ones carrying mutations in the candidate genes. The results showed no difference in MOSPD2 association for RHΔ*rop17* relative to wild type RH (Fig. 6A), indicating that ROP17 is not needed for MOSPD2 association with the PVM. Similarly, mutant parasites in *ROP1, ROP5, ROP18*, TGGT1_247350, and *GRA12* showed no change in host MOSPD2 association at the PVM relative to infection with RH-wild type (Fig. 6B-D). Together, these results suggest that while there are parasite proteins at the PVM that MOSPD2 is interacting with, at least the candidates tested so far are not individually responsible for the presence of MOSPD2 at this location.

**Fig 6:**
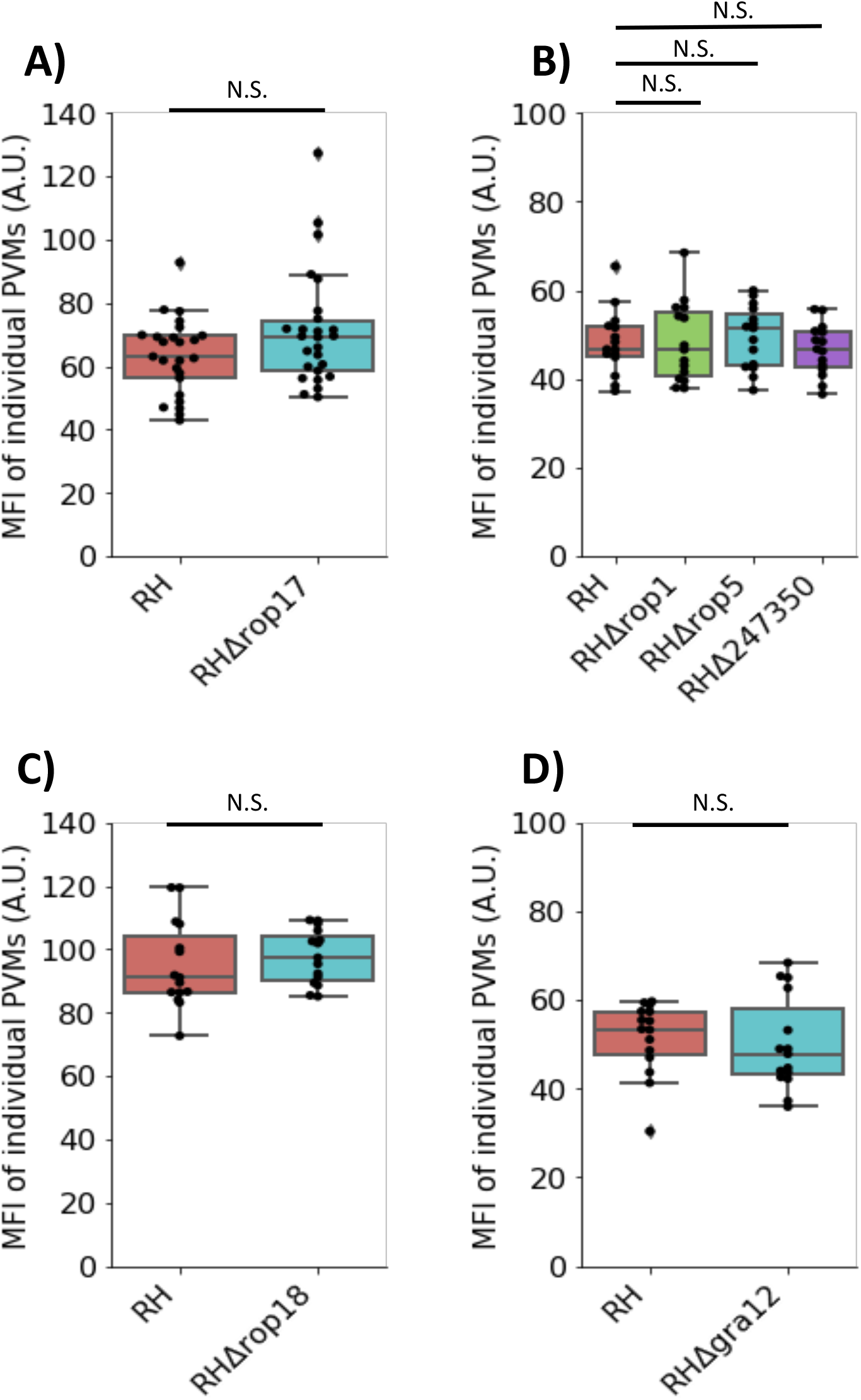
MOSPD2 association is not dependent on interacting proteins assessed so far. HFFs were infected with the indicated RH strains for 21-24 h. Monolayers were fixed with methanol then stained for endogenous MOSPD2. Fluorescence intensity of host MOSPD2 was quantified at the PVM using Fiji (see methods) for: A. RH, RHΔrop17, B. RH, RHΔrop1, RHΔrop5, RHΔ247350, C. RH, RHΔgra12, D. and RH, RHΔrop18. Data are from one experiment that pooled three biological replicates. One-way ANOVA was used to test significance in (B). N.S. indicates not statistically significant (P≥0.05). Student’s t test was used to test significance in (A), (C), and (D). All data are representative of three biological replicates pooled from one experiment. One of two fully independent experiments is shown.

The MOSPD2 association could either be pre-existing MOSPD2 recruited to the PVM-ER interface from other cellular compartments or newly synthesized material that is drawn to this location as its first (and only) destination. To distinguish between these competing hypotheses, HFFs were pre-treated with cycloheximide (CHX) for 1 hour then infected with ME49 for 6 additional hours (Fig. 7A). CHX blocks host cell translation and it was previously shown that *Toxoplasma* tachyzoites can grow for at least 16 hours in host cells blocked for protein translation [49]. Quantitation of the MOSPD2 fluorescent signal at the PVM in HFFs pretreated with CHX was greatly reduced relative to untreated controls (Fig. 7B and C). To determine if CHX was somehow disrupting the trafficking and insertion of parasite-derived PVM proteins, we infected and treated HFFs as described above and stained for GRA7 and MAF1b.

**Fig 7:**
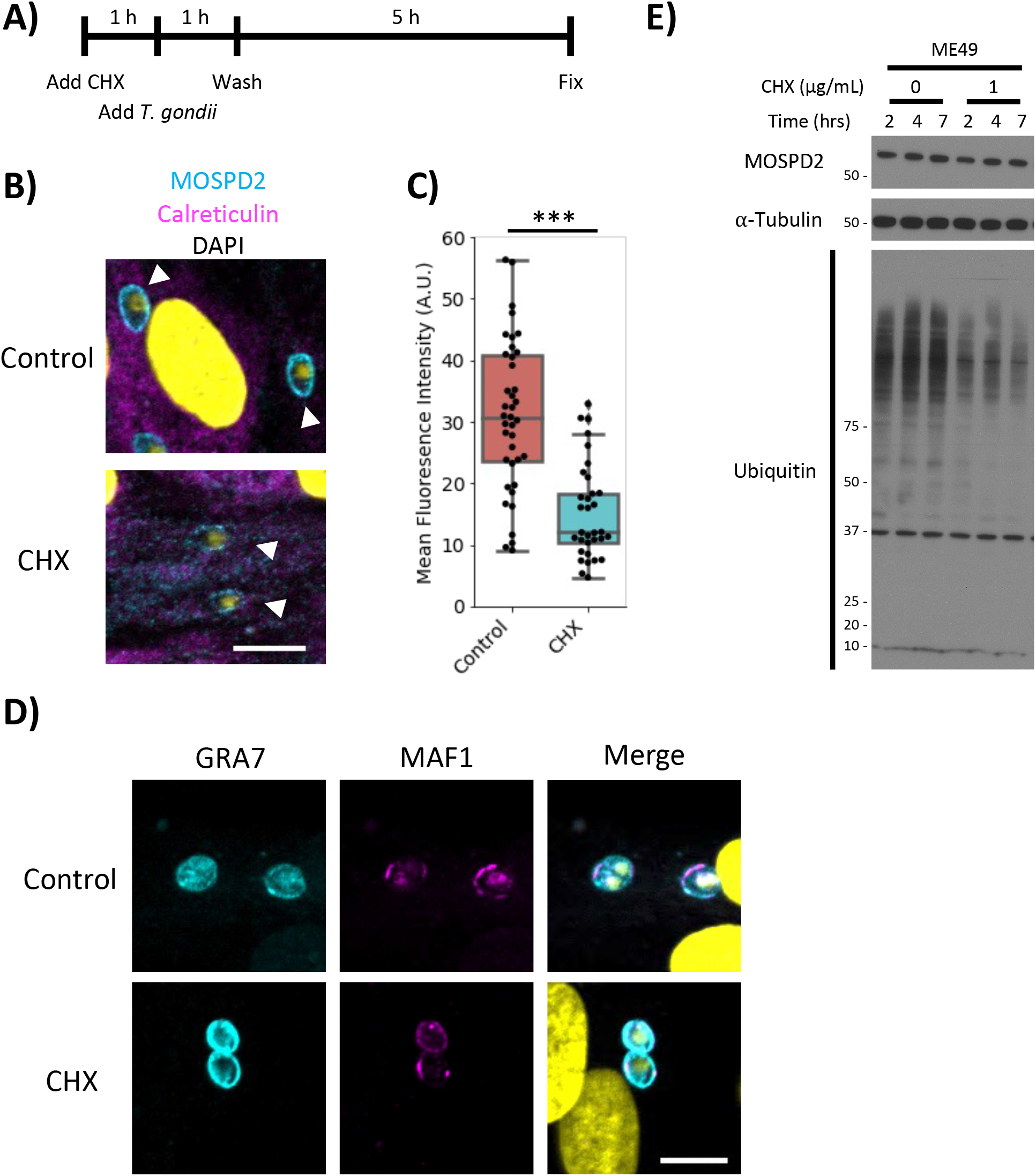
Host translation is necessary for prominent MOSPD2 association at the PVM. A. Schematic showing the addition of CHX (1 μg/mL) to HFFs 1 h prior to the addition of ME49. The monolayer was washed 1 h after the addition of the parasite. Media was replaced with either control or CHX-containing media where appropriate. B. Representative images of control and CHX-treated HFFs infected with ME49 for 6 h. Cell monolayers were fixed with methanol and stained with MOSPD2 (cyan) and calreticulin (magenta). DAPI is in yellow. Scale bar = 10 μm. C. Quantification of MOSPD2 signal at ME49 PVMs at 6 hpi from (B). Data are from one experiment that pooled 3 biological replicates. Data are representative of two fully independent experiments. Significance was tested using the Student’s t test (*** indicates P<0.001). D. Infected HFFs prepared as described in (A). Monolayers were stained for GRA7 (cyan), MAF1b (magenta), and DAPI (yellow). Data are representative of two experiments. Scale bar = 10 μm. E. Representative Western blot of whole cell lysate from ME49-infected HFFs prepared as in (A). The blot was probed with antibodies against host MOSPD2, α-Tubulin, and Ubiquitin. Data are representative of three independent experiments.

Imaging these monolayers showed GRA7 and MAF1b localized to the PVM, as expected (Fig. 7D). To determine whether global levels of MOSPD2 change with CHX treatment, a Western blot was performed under the same conditions as the immunofluorescence assay described above. Levels of MOSPD2 did not perceptibly change up to 7 hours post-CHX treatment, indicating MOSPD2 was not being degraded to any significant extent during this period of treatment (Fig. 7E). To confirm translation was successfully inhibited in HFFs, ubiquitin levels were assessed under the assumption that the proteasome would still be rapidly degrading ubiquitinated proteins during CHX treatment, a result reported in other cell types [50]. Western blot analysis of ubiquitin levels showed a marked reduction of ubiquitinated proteins after CHX treatment relative to untreated controls (Fig. 7E), indicating that the CHX treatment was working, as expected. Together, these results indicate that association of MOSPD2 with the PVM depends on active translation, suggesting that newly synthesized material is what associates with the PVM, rather than previously synthesized MOSPD2 being “stolen” from other membranes.

Three main domains have been previously described for MOSPD2: the CRAL/TRIO, MSP, and a c-terminal tail anchor (Fig. 8A). To determine which of these domains plays a role in association with the PVM, V5-tagged mutant constructs of each domain were generated and overexpressed in wild type HFFs using lentivirus transduction methods. For the CRAL/TRIO and tail anchor regions, the mutants were deletion constructs (ΔCT and ΔTA, respectively), whereas for the MSP domain, point mutations at Arg404 and Leu406 were generated (R404D/L406D) since these mutations were previously reported to destroy the ability of MSP to associate with the FFAT motifs they normally recognize [35]. Western blot analysis shows protein bands at the expected size for each mutant and in the wild type (Fig. 8C), albeit with different expression levels for each, possibly because of toxicity associated with some of these constructs. To determine if association at the PVM was still achieved with the mutant constructs, these HFF cell lines were infected with ME49 and stained by immunofluorescence. Association at the PVM was seen in infected cells expressing the wild type, ΔCT, and R404D/L406D constructs, although the association was reduced for the ΔCT version (Fig. 8B). For ΔTA, however, which was expressed at reduced but still detectable levels, no PVM association was seen (Fig. 8B). Quantifying the ratio of PVM-localized V5 signal (MOSPD2) relative to signal within the host cytosol showed a significant drop in the ΔCT line and no significant PVM-enrichment in the ΔTA line compared to the wild type control. Interestingly, the double point mutant appeared to have the opposite effect; i.e., a greater tendency to localize to the PVM than the wild type control (Fig. 8D). Together, these results indicate that the CRAL/TRIO and tail anchor of MOSPD2 are necessary for its efficient association at the *Toxoplasma* PVM whereas the MSP domain might even work against such association.

**Fig 8:**
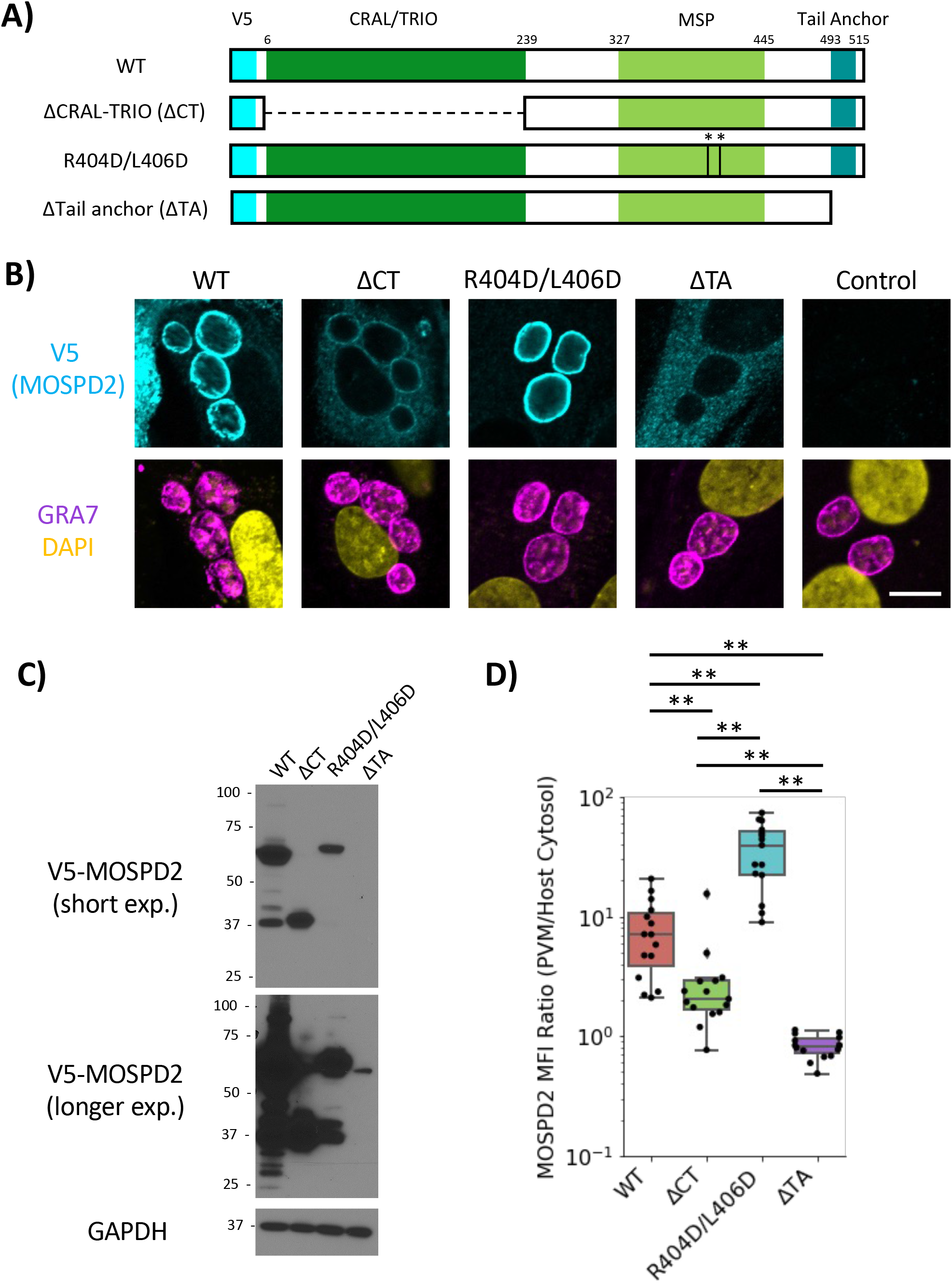
The tail anchor and CRAL/TRIO of MOSPD2 are necessary for association at the PVM. A. Schematic showing wild type MOSPD2 and mutant constructs ΔCRAL-TRIO (ΔCT), the double point-mutant R404D/L406D, and ΔTail-Anchor (ΔTA). V5 (cyan) is an inserted V5-epitope tag. B. Stable HFF cell lines over-expressing wild type, mutant MOSPD2 constructs outlined in (A), or control cells that did not receive lentivirus vector. HFFs were infected with ME49 for 21-24 h then fixed with methanol and probed for GRA7 (magenta), V5 (MOSPD2, cyan), and DAPI (yellow). Images are representative from 3 independent experiments. Scale bar = 10 μm. C. Uninfected whole cell lysates were lysed in RIPA. Western blot on the lysates was probed for V5 (MOSPD2), and GAPDH. Short and long exposure of the blot was used to capture ΔTA protein without saturating the other sample lanes. Data are representative of one experiment. D. Quantitation of V5 signal at the PVM as a ratio of PVM V5 signal divided by cytosolic signal. Data shown are from one experiment that pooled three biological replicates. One of three fully independent experiments is shown. Significance was tested using a One-way ANOVA and Tukey Post-HOC test (** indicates P<0.01).

Arginine-rich amphipathic helices (AHs) present in many ROPs are known to be sufficient to enable association with the PVM [26], and MOSPD2 was recently described to harbor one positively charged AH within the CRAL/TRIO [35]. Using AlphaFold [51], [52], MOSPD2 is also predicted to have a second c-terminal AH, proximal to the tail anchor. To determine if these structures are sufficient for PVM-association, the CRAL/TRIO AH (AH-CT) and the AH just preceding the tail anchor along with the tail anchor itself (AH-TA) were conjugated to the C-terminal end of eGFP and transiently transfected into HFFs (Fig. 9A). Live-cell imaging and quantitation of eGFP signal showed prominent association of eGFP-MOSPD2 with the PVM while the negative control had no apparent association (Fig. 9B and 9C). AH-CT and AH-TA constructs, like the negative control, also had no apparent association with the PVM (Fig. 9B and 9C). Together, these results indicate that neither the AH-CT nor the tail anchor and its associated AH are sufficient for association with the PVM.

**Fig 9:**
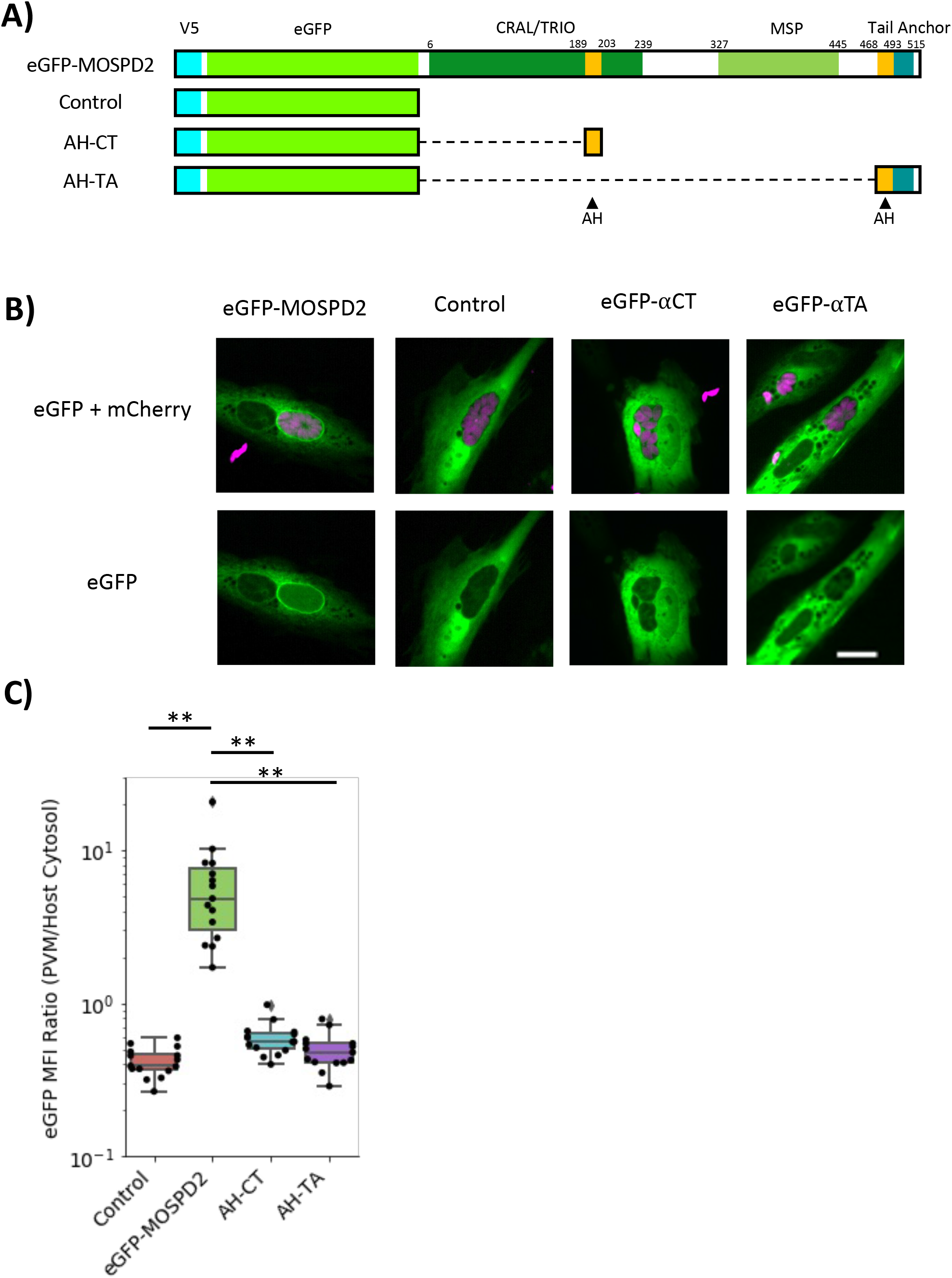
The CRAL/TRIO amphipathic helix or tail anchor are not sufficient to associate with the Toxoplasma PVM. A. Schematic showing eGFP constructs conjugated to a full length MOSPD2 (eGFP-MOSPD2), the amphipathic sequence in the CRAL/TRIO (AH-CT), the tail anchor including the proximal amphipathic sequence (AH-TA), and a control. AH = amphipathic helix. B. HFFs transiently transfected with the constructs from (A). HFFs were infected 7 h post-transfection with RH mCherry (magenta) for 21 h then imaged live. Images are show from 1 experiment and representative of at least 3 fully independent experiments. Scale bar = 20 μm. C. Quantitation of PVM localized eGFP signal normalized to host cytosol for each individual host cell. Significance was tested using a One-way ANOVA and Tukey Post-HOC test (** indicates P<0.01).

MOSPD2 has been reported to not be essential for human cell growth [53] and so to generate a stable knockout in HFFs, a CRISPR guide targeting MOSPD2 was transduced into host cells while control cells received a non-targeting (Control) guide. After puromycin-selection for uptake of the gRNA-containing constructs, which also carry a puromycin-resistance cassette, knockout (ΔMOSPD2) efficiency was assessed in the population by Western blot.

Immunostaining showed a band for endogenous MOSPD2 in the HFFs receiving no virus or the Control gRNA while no detectable signal was seen in the ΔMOSPD2 knockout population (Fig. 10A), indicating the knockout efficiency was extremely high. To further confirm the ΔMOSPD2 HFFs were knockouts, they were infected for 21-24 hours and stained for endogenous MOSPD2. *Toxoplasma* vacuoles in “No-virus” and Control HFFs had association of MOSPD2 while all observed ΔMOSPD2 HFFs had no detectable MOSPD2 at the PVM confirming efficient knockout (Fig. 10B). To determine whether loss of MOSPD2 has an impact on *in vitro* growth of *Toxoplasma* tachyzoites, plaque assays were performed in triplicate for each of Type I/II/III parasites in Control vs. ΔMOSPD2 HFFs. The results (Fig. 10C) showed no significant difference in plaque size after growth on the two host cell lines for Type I (RH) and Type II (ME49). For Type III (CTG), however, there was a small difference that showed statistical significance with smaller plaques in the ΔMOSPD2 vs. Control HFFs. Examination of the triplicate flasks contributing to this difference, however, revealed considerable variability in overall plaque size with just one of the three flasks containing Control HFFs having unusually large plaques and just one flask with the ΔMOSPD2 HFFs having unusually small plaques (Fig. 10D). Examining the nine pair-wise comparisons for these two sets of three replicate flasks shows only one difference is significant (that between the two flasks just mentioned). Overall, these data indicate that loss of host MOSPD2 has little, if any, impact on growth in HFFs *in vitro*.

**Fig 10:**
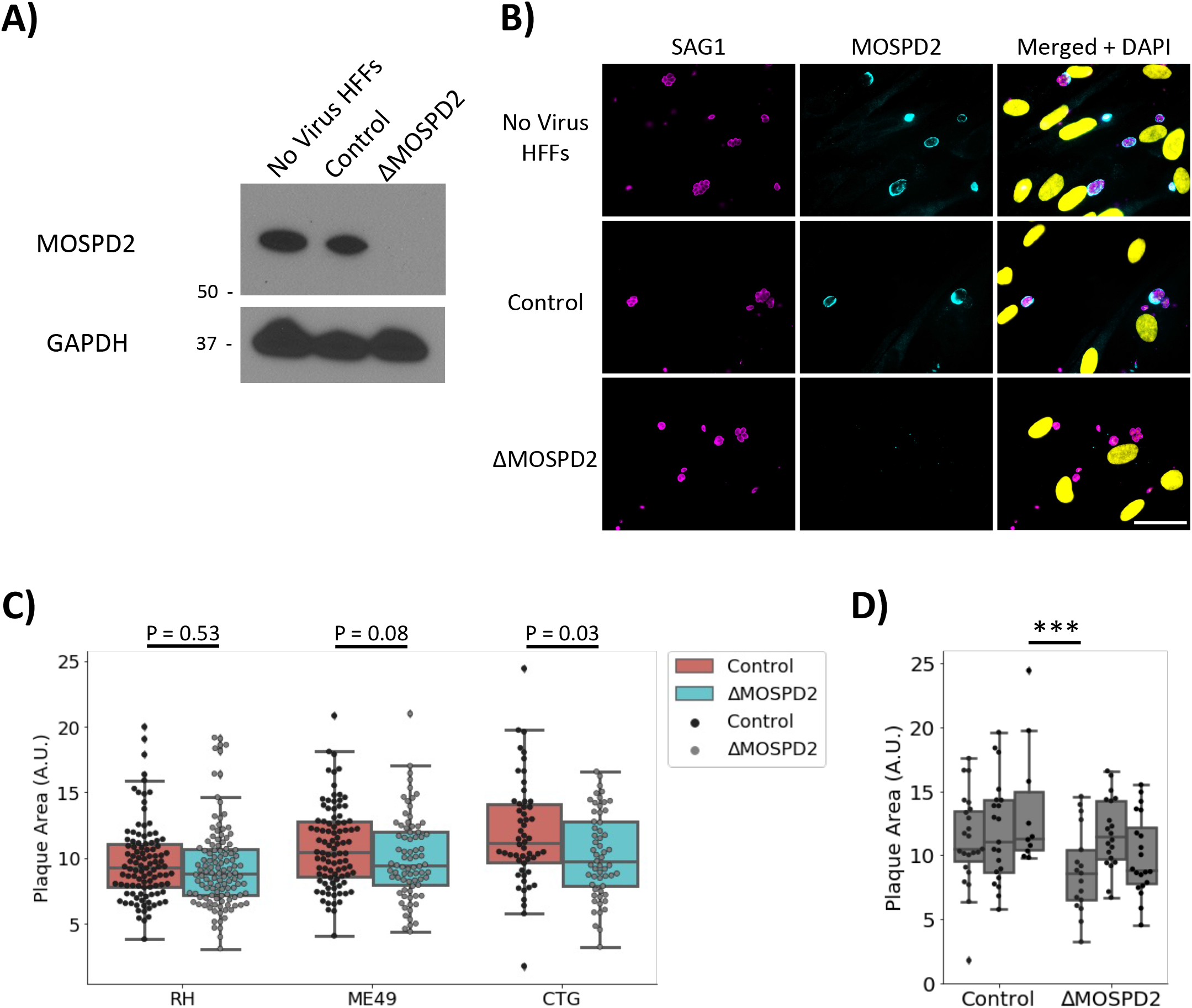
Ablation of MOSPD2 has little, if any, effect on Toxoplasma growth in vitro. A. HFF cell lines were generated using lentivirus to introduce CRISPR/Cas9 guides for MOSPD2 or a non-targeting control. Transduced HFFs were selected for using puromycin and expanded. Western blot for whole cell lysates from HFFs receiving no virus, Control (non-targeting control), and ΔMOSPD2 (MOSPD2 guide) was stained for endogenous MOSPD2 and GAPDH. B. No Virus HFFs, Control, and ΔMOSPD2 HFFs were infected with RH for 21-24 h, then fixed with methanol, and stained for SAG1 (magenta), MOSPD2 (cyan), DAPI (yellow). Scale bar = 40 μm. C. Control and ΔMOSPD2 cells were infected with RH, ME49, or CTG parasites for 10 days (RH), or 13 days (ME49, CTG). Monolayers were then stained with crystal violet and allowed to dry. Plaque area for each condition was measured in Fiji and plotted. Data are from one of two representative experiments each consisting of three biological replicates. Significance was tested using Student’s t test to do pair-wise comparisons. D. Plotted individual replicates from CTG in (C). Significance was tested using a One-way ANOVA and Tukey Post-HOC test (** indicates P<0.001).

The fact that MOSPD2 normally localizes to the ER and has a prominent role in mediating contact sites between host organelles suggested the possibility that it might have a role in ER association with the PVM in *Toxoplasma*-infected cells. To test this hypothesis, wild type and ΔMOSPD2 HFFs were infected with RH for 6 and 24 hours and then imaged using electron microscopy. Control HFFs showed strong HMA and host ER association with the PVM, as expected (Fig. 11A); interestingly, however, infected ΔMOSPD2 HFFs also exhibited clear association of the PVM with these two classes of host organelle (Fig. 11B). To determine if there were any changes in the fraction of PVM associated with ER, this was quantified in Control and ΔMOSPD2 HFFs. The results for both 6- and 24-hours post-infection showed no significant difference in ΔMOSPD2 relative to Control HFFs (Fig. 11C). These results argue against MOSPD2 playing an important role in mediating association of host ER with the *Toxoplasma* PVM.

**Figure 11:**
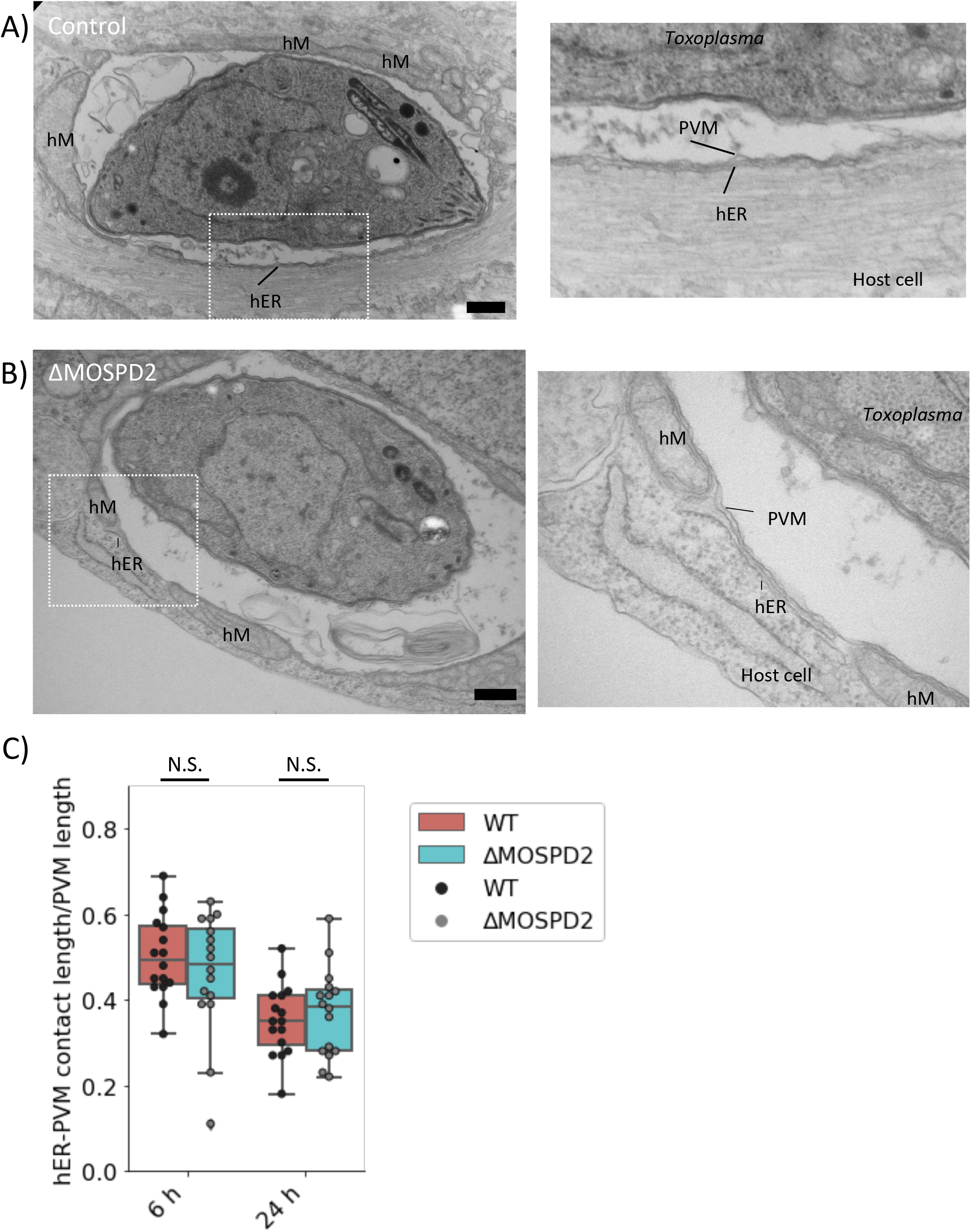
MOSPD2 ablation does not affect host ER association with the PVM. A. Left panel, electron micrograph images of RH-infected control HFFs 6 hpi. Right panel, zoomed image with labels marking Toxoplasma, PVM, host ER (hER), host Mitochondria (hM), and host cell. B. Left panel, image showing a RH-infected ΔMOSPD2 HFF 6 hpi. Right panel, zoomed image of the boxed region showing labeled Toxoplasma, hM, hER, host ribosomes, and the host cell. C. Quantitation of hER-PVM contact sites around the PVM at 6 and 24 hpi. Values are normalized to the circumference of the PVM and shown as a ratio of hER to PVM length. Significance was tested using Student’s t test to do pair-wise comparisons.

It has been hypothesized that the arginine-rich AH domain of ROPs drives association with the PVM through the attraction of the AH to the negative curvature of the PVM [26], but this possibility has not been directly tested. The MSP domain is known to be a protein-protein interactor [54] and so we decided to test whether the MSP domain might be interacting with the ROP-AH domain and whether this, in fact, is what draws ROP proteins to the PVM. To test this possibility, Control and ΔMOSPD2 HFFs were infected with ME49, and then fixed and stained with a monoclonal antibody that recognizes both ROP2 and ROP4 [55], or an antibody that recognizes the HA-tag on ROP17-3xHA (ROP2, ROP4 and ROP17 all have arginine-rich AH domains at their N-termini). The results showed no difference in association of ROP2/4 and ROP17 between the Control and the ΔMOSPD2 knockout cells (Fig. 12). Together with the results described above (Fig. 6), this indicates that association with the PVM of at least these three ROP proteins is neither responsible for, nor mediated by, MOSPD2.

**Fig 12:**
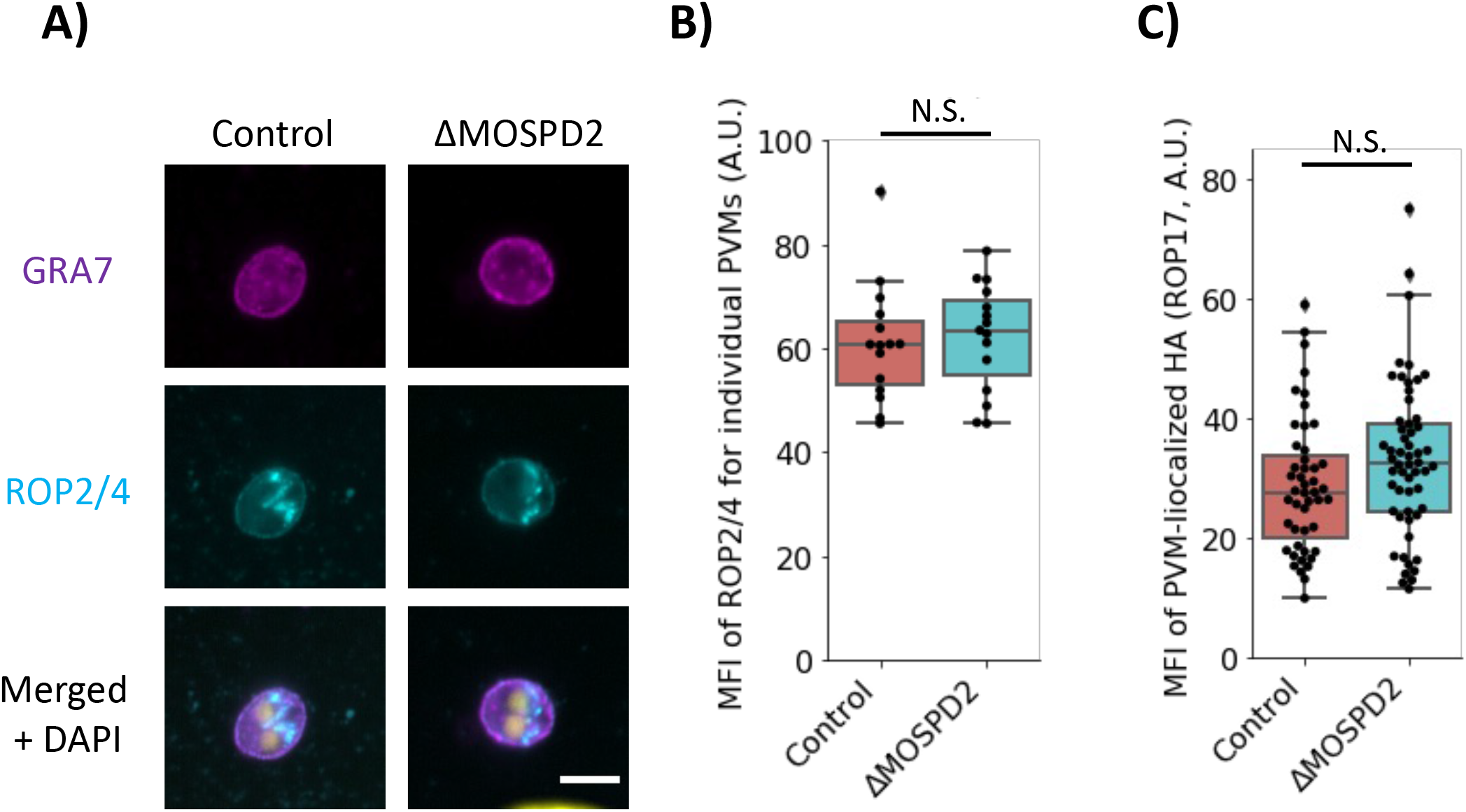
ROP2/4 and ROP17 association is not dependent on MOSPD2. A. Control and ΔMOSPD2 HFFs were infected with ME49 for 3 h and fixed with methanol. Monolayers were stained for ROP2/4 (cyan) and GRA7 (magenta). Fluorescence intensity of ROP2/4 signal was quantified for individual vacuoles in Fiji and MFI for individual vacuoles was plotted. Data are from one of two representative experiments each consisting of three biological replicates. Significance was tested using Student’s t test. Scale bar = 5 μm. B. Quantitation of sample data from (A). C. Control and ΔMOSPD2 HFFs were infected with RHΔrop17::ROP17-3xHA for 2-3 h then fixed with 4% formaldehyde. Samples were partially permeabilized using 0.02% Digitonin for 1-3 min, blocked with 3% BSA in PBS, and stained for HA (ROP17) and SAG1. Quantitative data shown are from one of two experiments each consisting of three biological replicates. Significance was tested using Student’s t test.

## Discussion

The PVM is a dynamic interface that allows *Toxoplasma* tachyzoites to co-opt host functions while remaining “hidden” from detection. Here, we dissected possible mechanisms and consequences of the dramatic presence of host MOSPD2 at this interface, a phenomenon that seems unlikely to be a chance event. One possible role for this association is that the CRAL/TRIO domain of MOSPD2 might bind to lipid substrates and move them between membranes, a function described for other proteins with similar domains [56]–[58]. Thus, MOSPD2-PVM association could be one of the ways *Toxoplasma* acquires lipids from its host, including lipids needed to expand the PVM and elaborate the IVN as the parasites grow. Future work assessing lipid scavenging in wild type and MOSPD2 knockout cells will be needed to address this possibility.

Another possibility is that MOSPD2 association at the PVM could be related to the immune response and so its function will only be revealed when infection of a cell type other than HFFs is examined or when cells are stimulated by cytokines like IFN-γ. For example, monocytes and macrophages have previously been shown to express higher levels of MOSPD2 [59]. The migration of monocytes has also been shown to be dependent on this host protein [59]. Our studies, however, have all been in nonmotile HFFs, precluding an assessment of the role of such enrichment in cell migration. *Toxoplasma* is known to induce hypermigratory phenotypes in monocytes, macrophages, and dendritic cells [60]–[65]. One *Toxoplasma* protein involved in this hypermigration is ROP17 [64], a protein specifically enriched in the MOSPD2 immunoprecipitation. It is therefore appealing to hypothesize that the association of MOSPD2 at the PVM could be a mechanism by which *Toxoplasma* influences monocyte migration. Future work could focus on detailing whether MOSPD2 association in monocytes impacts cell migration upon infection with *Toxoplasma*. Ultimately, however, *in vivo* studies will be needed and while viable MOSPD2 knockout mice have been reported [53], discriminating between direct and indirect effects of such a disruption will be extremely difficult; i.e., determining whether any difference in the course of an infection with *Toxoplasma* in such animals is because of a specific role of MOSPD2 at the PVM or is due to a generalized effect on the host’s overall metabolism, immunity or other characteristic.

In terms of the mechanism of MOSPD2-PVM association, we found that host mitochondria at the PVM preclude MOSPD2 association with this membrane. This seems most likely to relate to the fact that MOSPD2 is typically found embedded within the ER and, in any one patch of PVM at any one time, direct association can be with either host mitochondria or ER, but not both. Further work will be needed to determine whether other PVM-associating proteins show similar specificity in their location and, therefore, whether the mutually exclusive association of host mitochondria and MOSPD2 is mediated by the bulk size of mitochondria and ER or is due to more molecule-level effects such as localized differences in the lipid composition of the PV membrane. It is known, for example, that HMA facilitates lipid scavenging [66] and a possible hypothesis would be that the PVM’s lipid composition or some other aspect of the PVM is altered when mitochondria associate with the vacuole. Although studies have investigated lipids important for *Toxoplasma*’s growth (reviewed in [67]), characterization of the PVM has been difficult in the context of lipids due to the fragile nature of the vacuole and arduous task of separating host and parasite membranes/lipids.

We show that there are substantial differences in the extent of MOSPD2 association depending on the parasite strain used to infect HFFs. Many polymorphisms are known to exist between Types I/II/III parasites, including in expression levels and/or sequence of ROP5, ROP17 and ROP18 [29]–[31]. All of these co-precipitated with MOSPD2 from ME49-infected cells but none appeared individually responsible for MOSPD2’s association with the PVM. Fortunately, F1 progeny from crosses between Type I/II/III strains exist [68] and have been mapped. Therefore, phenotyping them with respect to MOSPD2 association should reveal the locus or loci involved, even if multiple such loci contribute collectively to the strain-specific difference; i.e., even if it is a quantitative trait.

The MOSPD2-*Toxoplasma* interactome at the PVM identified additional parasite proteins known to be at this host-parasite interface [20], [25], [30], [44]–[48]. Among these, ROP1 and the polymorphic ROP5 and ROP17 are of particular interest due to the presence of a short linear motif that resembles a FFAT (two phenylalanine in an acidic tract) [69]; such a motif in other host proteins is known to bind to the MOSPD2 MSP domain allowing MCS to occur between this ER-resident protein and other organelles [35]. Although we found that deletion of any one of these *ROP* loci on their own did not affect MOSPD2 recruitment, the possibility certainly remains that these and related ROP proteins collectively mediate MOSPD2 association with the PVM and that removal of multiple such proteins would markedly reduce MOSPD2 association. Alternatively, these parasite PVM proteins might associate with MOSPD2 indirectly, perhaps through interactions with other molecules, for example with specific lipid domains, as discussed above.

Mutant constructs in each of the three major domains of MOSPD2 revealed that the CRAL/TRIO and tail anchor play a role in, but are not sufficient for, MOSPD2’s localization to the PVM. This strongly suggests that MOSPD2 must be anchored into a membrane for it to be enriched at the PVM, but which membrane, the PVM-proximal side of host ER or the PVM itself, is not addressed by our data. CHX treatment showed that translation is necessary for association suggesting that the MOSPD2 accumulating at the PVM is newly translated rather than being taken from pre-existing supplies in the cell, although we cannot exclude the possibility that inhibiting translation eliminates a short-lived host protein necessary for such accumulation. For example, tail-anchored proteins are known to be chaperoned by a class of proteins in the GET pathway in yeast and TRC pathway in mammals once their translation is completed [70]. Perhaps one or many of these chaperones is short-lived, disrupting the pathway of tail anchor insertion into membranes. Certainly, a GET/TRC seems likely to be involved in the overall process but whether such is of parasite or host origin is not easily predicted. If it is of parasite origin, it could be polymorphic and explain the strain-specific difference. The *Toxoplasma* genome encodes strongly predicted GET/TRC proteins [ToxoDB.org] but there is no evidence reported that any of these are secreted outside the parasite, suggesting that they instead perform the usual duties for such proteins within the parasite (which are known to have their own tail-anchored proteins [71], [72] and so in need of their own GET/TRCs). Whether GET/TRCs are involved in MOSPD2’s association with the PVM, and whatever their source, this does not address which membrane, PVM or ER, harbors MOSPD2’s tail-anchor and the immune-electron microscopy performed here does not have the resolution needed to address this crucial point. Future studies could aim at characterizing this using a combination of molecular, genetic, and structural biology techniques.

The extent of the association of MOSPD2 with the PVM is, so far, one of the greatest of any host protein studied to date in both the degree and specificity of the association. It has the potential, therefore, to reveal both important biochemistry and biology about the host-parasite interaction. The work described here reveals several important details about this phenomenon but much more work will be needed to reveal exactly how and why it is drawn to this site so strongly.

## Materials and Methods

### Parasite strains, culture, and infections

*Toxoplasma* tachyzoites used in this study include type I RH strain, type II ME49, and type III CTG. RH *Toxoplasma* lines include: RH::*MYR1*-3xHA [38], RHΔ*myr1* [37], RHΔhptΔku80 [73], RHΔ*rop17* [43]. ME49 *Toxoplasma* lines include: ME49Δ*hpt*, ME49Δ*myr1* [37], ME49 MAF1b-NHA [74]. CTG *Toxoplasma* line was CTG (also referred to as CEP) expressing GFP luciferase [75]. *Neospora caninum* was NC-1 [76].

These tachyzoites and all subsequently generated lines were propagated in human foreskin fibroblasts (HFFs) cultured in complete Dulbecco’s modified Eagle medium (DMEM) supplemented with 10% heat-inactivated fetal bovine serum (FBS; HyClone, Logan, UT), 2mM L-glutamine, 100 U/ml penicillin, and 100 μg/ml streptomycin at 37°C with 5% CO2. The HFFs were obtained from the neonatal clinic at Stanford University following routine circumcisions that are performed at the request of the parents for cultural, health, or other personal medical reasons (i.e., not in any way related to research). These foreskins, which would otherwise be discarded, are fully deidentified and therefore do not constitute human subjects research. To obtain parasites, infected monolayers were scraped and the host cells lysed by passage through a 25-gauge needle and counted using a hemocytometer.

### Cycloheximide (CHX) treatment

HFF monolayers were seeded on sterile, glass-coverslips 18 or more hours prior to adding 1 μg/mL or 10 μg/mL cycloheximide (CHX, Millipore Sigma). HFFs in 24-well or 6-well dishes were incubated with CHX for 1 h before adding ME49 and pulse spun at 82 g for 1 second to help the parasites make contact with the host cells. Parasites were cultured on HFFs for 1 h then washed, and control or CHX-containing media was added where appropriate. The infected monolayers were then incubated for 5 additional hours prior to carrying out IFA protocols.

### Immunofluorescence assay (IFA)

Infected cells grown on glass coverslips were fixed and permeabilized using 100% cold methanol for 15 min. Samples were washed 3 times with phosphate-buffered saline (PBS) and blocked using 3% bovine serum albumin (BSA) in PBS for 30 min at room temperature (RT). HA was detected with rat monoclonal anti-HA antibody 3F10 (Roche), SAG1 was detected with mouse anti-SAG1 monoclonal antibody DG52 [77], GRA7 was detected with rabbit anti-GRA7 antibodies [78], V5 was detected with a mouse anti-V5 tag monoclonal antibody (Invitrogen), MOSPD2 was detected with a rabbit polyclonal antibody (Millipore Sigma), Calreticulin was detected with a mouse monoclonal antibody FMC 75 (Abcam), ROP2/4 was detected with a mouse monoclonal antibody [55]. Primary antibodies were detected with goat polyclonal Alexa Fluor-conjugated secondary antibodies (Invitrogen). Both the primary and secondary antibodies were diluted in 3% BSA in PBS. Coverslips were incubated with the primary antibodies for 1 h at RT, washed, and incubated with secondary antibodies for 45 min. at RT. Vectashield with DAPI (4’,6-diamidino-2-phenylindole) stain (Vector Laboratories) was used to mount the coverslips on slides. Fluorescence was detected using wide-field epifluorescence microscopy or by confocal microscopy on an LSM 700 laser scanning confocal microscope. Images were analyzed using ImageJ software and the intensity levels of the images adjusted such that no data were removed from images. All images shown for any given condition/staining in any given comparison/data set were obtained using identical parameters unless otherwise stated.

For mitochondrial staining, HFF monolayers were infected as described. MitoTracker (Invitrogen) was used at manufacture’s recommendation prior to fixing with 4% formaldehyde in DMEM lacking phenol red for 15 minutes at room temperature. Samples were then washed and permeabilized with 0.2% Triton X-100 for 15 minutes at room temperature. IFA staining was done as described above.

### Partial Permeabilization

Parasites were syringe released as described and used to infect HFFs for 2-3 h, at which time the cells were washed with PBS and then fixed with 4% formaldehyde at room temperature for 15 min. Formaldehyde-fixed samples were rinsed once with PBS, permeabilized with 0.02% digitonin solution for 1-3 min, and then blocked with 3% BSA in PBS for 1 h at RT. Staining was performed as described above.

### IFA analysis

Quantification of MOSPD2 fluorescence around the PVM was performed in Fiji. To quantify fluorescence intensity, a polygonal region of interest (roi) was generated around the outside and inside of the PVM as defined by calreticulin or SAG1 staining to exclude the host cytosol, parasites and lumen of the PV. The roi was copied onto the MOSPD2 channel and pixel intensity was measured. Average fluorescence intensity for the roi was plotted as arbitrary units.

Alternatively, a single line 1 or 5 pixels wide was generated around the PVM, again guided by calreticulin/SAG1 staining or intra-parasitic fluorescent signal. It was copied onto the MOSPD2 channel and pixel intensity measured at each point on the line. To normalize across vacuoles, each pixel intensity value was divided by the median or mean pixel intensity for that given vacuole and the ratios generated plotted.

### Plasmid construction

For lentiviral plasmid constructs, standard molecular cloning techniques were used to amplify wild type MOSPD2 from a pCDNA construct (primers A1, A2, A20 and A21, [34]; a list of all primers used in this study can be found in Supplementary Table 1).

The amplified V5-MOSPD2 was subsequently inserted into pLenti-CMV-puro (Addgene plasmid no. 17452). MOSPD2 CRISPR guides for MOSPD2 (A3 and A4, [79]) and a non-targeting guide (A5 and A6, [79]) were annealed and ligated into lentiCRISPRv2 (a generous gift from Jan Carette, Stanford University, Addgene, plasmid no. 52961). Plasmids were subsequently sequenced to confirm correct inserts A22 and 3CMVFW (Sequetech.com, Sequetech Corporations).

For mutant MOSPD2 constructs, ΔCT (primers A9 and A10), ΔTA (primers A11 and A12), and RL/DD (primers A7 and A8) were generated using wild type pCDNA and ligated. Mutant constructs were then amplified (ΔCT and RL/DD: primers A1 and A2; ΔTA: primers A1 and A19) and inserted into pLenti DEST.

For eGFP constructs, the backbone originated from pCDNA-MOSPD2 [34]. The backbone for AH-TA was amplified using A9 and A25 from pCDNA-MOSPD2 [34]. The eGFP insert was amplified from CAS9_sgRNA [80] using primers A23 and A24. The negative control and AH-CT plasmids were constructed using AH-TA as a template and primers A11 and A28 then A26 and A27, respectively. The full length eGFP-MOSPD2 plasmid was constructed using primers A11 and A30 from HA-TA. The MOSPD2 insert was amplified using primers A29 and A31 from pCDNA-MOSPD2 [34]. Plasmids were ligated using standard molecular biology techniques.

### Mammalian cell culture and stable cell line generation

All mammalian cell lines were propagated in complete Dulbecco’s modified Eagle medium (DMEM) supplemented with 10% heat-inactivated FBS, 2 mM L-glutamine, 100 U/ml penicillin, and 100 mg/ml streptomycin at 37°C with 5% CO_2_, unless otherwise noted.

For preparation of lentiviruses, HEK 293T cells in 10-cm dishes were transfected at 80% confluence with the lentiviral plasmid pLenti-CMV-Puro (Addgene plasmid no. 17452) containing the gene of interest (2 mg) and the lentiviral packaging plasmids pVSV-G, pDVPR, and pAdvant (gifts from Jan Carette, Stanford University) using FuGENE HD transfection reagent (Promega) according to manufacturer’s instructions. After about 24 h, the medium was replaced with fresh medium. Approximately 48 h after transfection, the cell medium containing the lentivirus was harvested and filtered through a 0.45-μm filter and supplemented with 8 mg/ml protamine sulfate. To generate stable lines, HFFs in T25 culture flasks were then infected with the virus-containing medium (1-2.5 ml). The following day, the viral-containing medium was removed and replaced with fresh, antibiotic-free medium. The HFFs were allowed to recover for 48 h and then selected with medium containing 2 μg/ml puromycin for 3 days.

### Transient mammalian transfections

HFFs were grown on glass coverslips to ∼80% confluence and subsequently transfected with Lipofectamine LTX reagent (Invitrogen) and 500 ng of each pCDNA plasmid (with the tagged gene of interest as described above) according to the manufacturer’s instructions in antibiotic-free medium. Cells were incubated with the transfection reagent for ∼6-7 h and tachyzoites were added for another 21 h before imaging or fixing.

### *Toxoplasma* Transfections

All transfections were performed using the Amaxa 4D Nucleofector (Lonza) model. Tachyzoites were mechanically released in PBS, pelleted, and resuspended in 20 μl P3 primary cell Nucleofector solution (Lonza) with 5 to 20 μg DNA for transfection. After transfection, parasites were allowed to infect HFFs in DMEM.

### Gene disruption in *Toxoplasma*

A list of all sgRNA sequences and primers used in this study can be found in Supplemental Table 1. For gene disruption plasmids, guide RNAs, designed against a PAM site of each gene of interest, were cloned into the pU6-Universal plasmid (Addgene plasmid number 52694; http://n2t.net/addgene:52694; RRID:Addgene_52694). Parasites were transfected with the pU6-sgRNA plasmid containing the guide for GRA12 (A13 and A14) or TGGT1_247350 (A32 and A33) and allowed to infect HFFs in DMEM. Linear PCR-amplified hypoxanthine-guanine phosphoribosyl transferase (HPT) (primers A15 and A16 from pTKO2 [38]) was co-transfected with the GRA12 guide. After at least 18 hours of recovery time, transfected cell cultures were drug selected for 8 days with 25 μg/ml mycophenolic acid (MPA) and 50 μg/ml xanthine (XAN). Single clones were selected from the transfected populations in 96-well plates using limiting dilution in MPA/XAN-supplemented medium and PCR verified for gene disruption (A17 and A18).

### Western blotting

Cell lysates were prepared in Laemmli sample buffer (Bio-Rad) at the time points post-infection indicated. The samples were boiled for 5 min, separated by SDS-PAGE, and transferred to polyvinylidene difluoride (PVDF) membranes. The membranes were blocked with 5% nonfat dry milk or 5% BSA in Tris-buffered saline supplemented with 0.5% Tween 20, and proteins were detected by incubation with primary antibodies diluted in blocking buffer, followed by incubation with secondary antibodies (raised in goat against the appropriate species) conjugated to horseradish peroxidase (HRP) and diluted in blocking buffer. HA was detected using an HRP-conjugated HA antibody (catalog no. 12013819001; Roche), SAG2A was detected using rabbit polyclonal anti-SAG2A antibodies, VAP-A was detected using mouse monoclonal 4C12 (Santa Cruz), Calreticulin was detected with a mouse monoclonal antibody FMC 75 (Abcam), MOSPD2 was detected with a rabbit polyclonal antibody (Millipore Sigma), Ubiquitin was detected using rabbit polyclonal antibody (Thermo), α-Tubulin was detected using mouse monoclonal antibody B-5-1-2 (Signa-Aldrich), and GAPDH (glyceraldehyde-3-phosphate dehydrogenase) was detected using mouse monoclonal anti-GAPDH antibody 6C5 (Calbiochem). Horseradish peroxidase (HRP) was detected using an enhanced chemiluminescence (ECL) kit (Pierce).

### Plaque assay

Parasites were syringe released from HFFs and added to confluent HFFs in T25 flasks. After 10 or 13 days of undisturbed incubation at 37°C as described above, the infected monolayers were washed with PBS, fixed with methanol, and stained with crystal violet. The plaque area was measured in arbitrary units using ImageJ software.

### Transmission electron microscopy (TEM)

For ultrastructural observations of Toxoplasma-infected HFF for 24 h by thin section, samples were fixed in 2.5% glutaraldehyde in 0.1 mM sodium cacodylate and processed as described previously [81]. Ultrathin sections of infected cells were stained with osmium tetroxide before examination with Hitachi 7600 EM under 80 kV equipped with a dual AMT CCD camera system. Quantitative measurement of length for the PV membrane and host ER elements attached to this membrane using ImageJ was performed on 18 representative electron micrographs at low magnification to ensure the entire PV fit into the field of view.

### Immunoelectron microscopy (IEM)

Monolayers of HFF infected with Toxoplasma from 6 or 24 h were fixed in 4% paraformaldehyde (PFA; Electron Microscopy Sciences, PA) in 0.25 M HEPES (ph7.4) for 1 h at room temperature, then in 8% PFA in the same buffer overnight at 4°C. Samples were infiltrated, frozen and sectioned as previously described [82]. The sections were immunolabeled with rabbit anti-MOSPD2 antibody at 1/50 diluted in PBS/1% fish skin gelatin. The sections were then incubated with IgG antibodies, followed directly by 10 nm protein A-gold particles before examination with the EM.

### IPs for mass spectrometry

Immunoprecipitations (IP) to identify MOSPD2-interacting proteins in HFFs were performed as follows. One 10-cm dish of HFFs for each infection condition were seeded with 3 million HFFs 18-24 hours prior to infection. HFFs were infected with 16 million ME49Δ*hpt* parasites for 19 h. Infected cells were washed 3 times in cold PBS and then scraped into 1 ml cold cell lysis buffer (50 mM Tris [pH 8.0], 150 mM NaCl, 0.1% [vol/vol] Nonidet P-40 alternative [CAS no. 9016-45-9]) supplemented with complete protease inhibitor cocktail (cOmplete, EDTA free; Roche) and phosphatase inhibitor (PhosSTOP, Roche). Cell lysates were passed 3 times through a 25-gauge needle, followed by passage 3 times through a 27-gauge needle. After lysing, samples were incubated on ice for 30 additional minutes. The cell lysates were spun at 10,000 g for 10 min at 4°C to remove insoluble material and unlysed cells. Protein concentration was calculated using a Bradford Assay (Thermo Scientific). Equal amounts of protein (3,500 μg) from lysates were added to 100 μl magnetic beads conjugated to anti-V5 nanobodies (Chromotech), and the mixture was incubated rotating at 4°C for 1 h. Unbound protein lysate was removed, and the anti-V5 magnetic beads were then washed 10 times in cell lysis buffer containing protease inhibitor and lacking NP-40. V5-tagged MOSPD2 and associated proteins bound to the beads were delivered to the Stanford University Mass Spectrometry core for on-bead digestion.

### Mass spectrometry sample preparation

For on-bead digestion, beads were resuspended in 100 mM TEAB and mixed head-over-head on a ThermoLyne LabQuake shaker for 10 minutes. DTT was added to a final concentration of 10 mM at 55°C for five minutes followed by head-over-head mixing, at room temperature, for 25 minutes. Acrylamide was added at 30 mM for cysteine capping during head-over-head mixing for an additional 30 minutes. Trypsin/LysC (500 ng) was added for proteolysis overnight, at 37°C. Samples were then quenched with 5 uL of 50% formic acid, separated from beads, and cleaned by C18.

### Mass spectrometry

For peptides from on-bead digests, the samples were analyzed either on a Orbitrap Fusion tribrid mass spectrometer (Thermo Scientific) RRID:SCR_018702 or an Orbitrap Eclipse tribrid mass spectrometer (Thermo Scientific) RRID:SCR_022212, in both cases coupled to a Acquity M-Class liquid chromatograph (Waters Corporation). In brief, peptides were injected at a flow rate of 300 nL/min with a mobile phase A of 0.2% aqueous formic acid and a mobile phase B of 0.2% formic acid in acetonitrile. Peptides were directly injected onto a ∼25 cm in-house pulled-and-packed fused silica column with an I.D. of 100 microns. The column was packed with 1.8 micron C18 stationary phase, and the gradient was a 2-45% B, followed by a high B wash for a total gradient time of 180 min. The mass spectrometer was operated in a data dependent fashion using CID fragmentation in the ion trap to generate MS/MS spectra and HCD in the orbitrap following synchronous precursor selection for detection and quantification of report ions in the MS3 step.

### Mass spectrometric analysis

For a typical data analysis, Peptide spectra assignments and protein inferences were performed using Byonic v.4.2.4 (Protein Metrics), assuming fully-specific tryptic digestion and up to 2 missed cleavages, as well as common modifications such as methionine oxidation and cysteine alkylation.

## Data availability

The mass spectrometry proteomics data have been deposited to the ProteomeXchange Consortium via the PRIDE [83] partner repository with the dataset identifier PXD038158 and 10.6019/PXD038158.

## Acknowledgments

We thank all members of our laboratory for helpful comments and input to the experiments and manuscript and Melanie Espiritu for help with tissue culture and ordering. We also thank Ryan Leib, Fang Liu, Kratika Singhal, and Norah Brown for their support at the Stanford SUMS core. We would like to thank the Bogyo lab for allowing us to use their confocal microscope. We thank Christine Peters from the Carette lab for providing input and gifting reagents for knocking out MOSPD2. We thank the excellent technical staff of the Electron Microscopy Core Facility at the Johns Hopkins University School of Medicine and at the Yale University School of Medicine.

This project has been funded in whole or part with: federal funds from the National Institute of Allergy and Infectious Diseases, National Institutes of Health, Department of Health and Human Services under Award Numbers NIH RO1-AI21423 (JCB), NIH RO1-AI129529 (JCB); 5T32AI007328-30 to AF; Gilliam Fellowship HHMI to AF. Mass spectrometry measurements were performed at the Vincent Coates Foundation Mass Spectrometry Laboratory, Stanford University Mass Spectrometry (SUMS - RRID:SCR_017801). This work was supported in part by NIH P30 CA124435 utilizing the Stanford Cancer Institute Proteomics/Mass Spectrometry Shared Resource. This work includes data collected using the Orbitrap Eclipse mass spectrometer system (RRID:022212) which was purchased using funding through the National Institutes of Health Shared Instrumentation Grant 1S10OD030473-01. The funders had no role in study design, data collection and analysis, decision to publish, or preparation of the manuscript.

---

**Supplementary Table 1:**
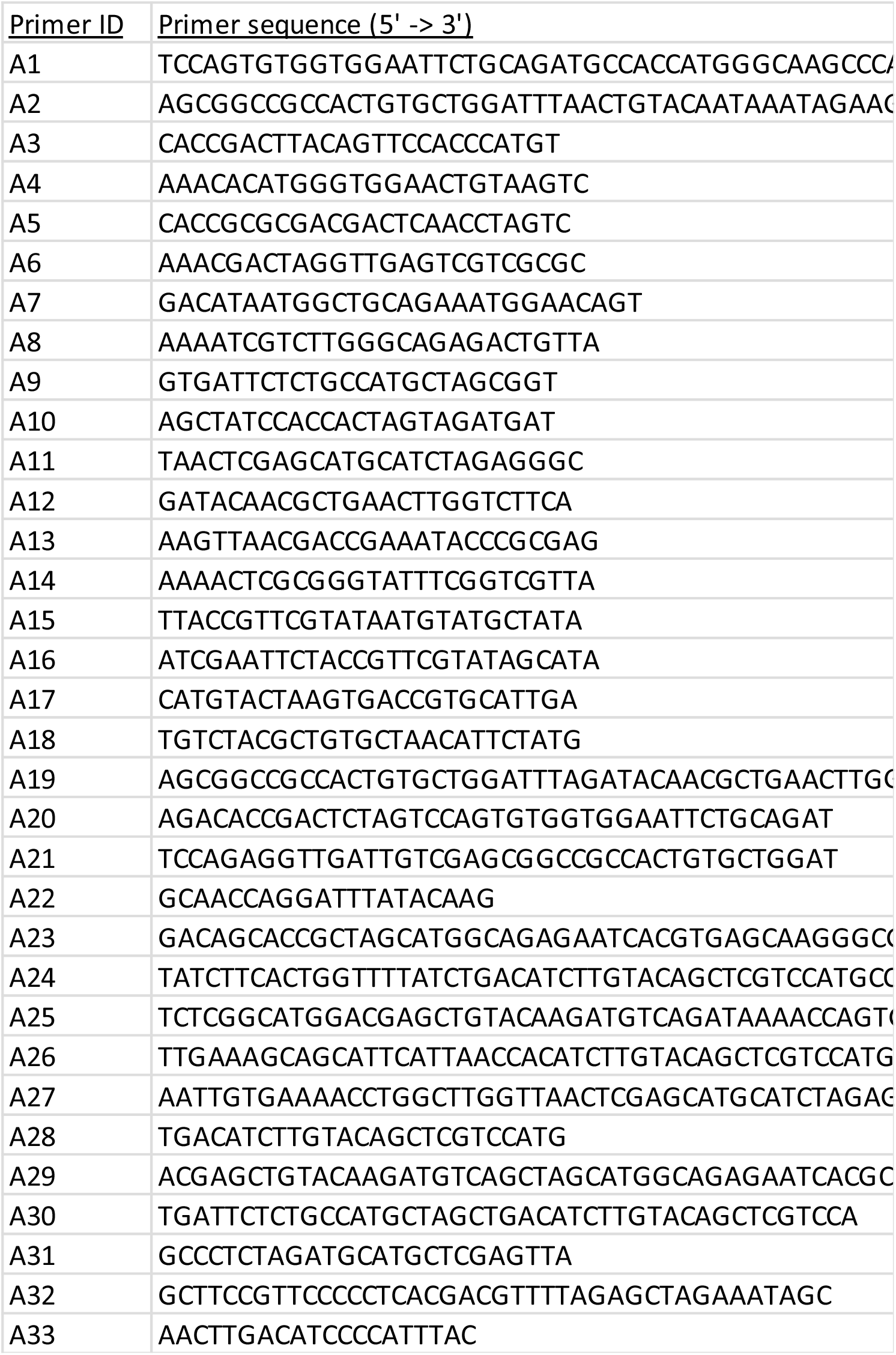
Primers

## Supplementary Data Set 1: V5-MOSPD2 IP Ranked protein list

Complete mass spectrometry data from Human and *T. gondii*.

